# LexicMap: efficient sequence alignment against millions of prokaryotic genomes

**DOI:** 10.1101/2024.08.30.610459

**Authors:** Wei Shen, John A. Lees, Zamin Iqbal

## Abstract

Alignment against a database of genomes is a fundamental operation in bioinformatics, popularized by BLAST. However, the rate at which microbial genomes are sequenced has continued to increase, and there are now datasets in the millions, far beyond the abilities of existing alignment tools. We introduce LexicMap, a nucleotide sequence alignment tool for efficiently querying moderate length sequences (>250 bp) such as a gene, plasmid or long read against up to millions of prokaryotic genomes. A key innovation is to construct a small set of probe *k*-mers (e.g. n = 20,000) which are selected to efficiently sample the entire database to be indexed, such that every 250 bp window of each database genome contains multiple seed *k*-mers each with a shared prefix with one of the probes. Storing these seeds in a hierarchical index enables fast and low-memory alignment. We benchmark both accuracy as the query diverges from the match in the database, and potential to scale to databases of millions of bacterial genomes, showing that LexicMap achieves comparable accuracy with state-of-the-art, but with greater speed and lower memory use. We then benchmark LexicMap on small/diverse (GTDB) and large/redundant (AllTheBacteria and GenBank+RefSeq) databases. Alignment of a single gene against 2.34 million prokaryotic genomes from GenBank and RefSeq takes 3 (rare gene) to 33 (16S rRNA gene) minutes. Full alignment against all bacterial genomes is now possible in minutes with modest resources, supporting querying at scale which will be useful for many biological applications across epidemiology, ecology and evolution.

LexicMap produces output in standard formats including that of BLAST and is available under MIT license at https://github.com/shenwei356/LexicMap.

## Introduction

Alignment of sequences to a single reference genome is a well-studied problem^1–4^. For specified gap and mismatch costs, dynamic programming is guaranteed to obtain the optimal solution^5–7^, but processing time scales as the produce of the reference genome size and the query size. This is too slow in practice, so the challenge is to find faster shortcuts that ideally are still guaranteed to give the optimal solution. Rapid progress has recently been made on this front^8–12^.

The problem we set out to address is that of aligning to a database of genomes from across the bacterial phylogeny, as popularised by BLAST^1^. As the amount of publicly available bacterial sequence has grown over recent years, the proportion of bacterial genomes which web BLAST is able to search has dropped exponentially^13^. The primary use cases of alignment to either all bacterial genomes or a representative set are: determining where a specific sequence has been seen before, finding the host range of a mobile element or gene, or locating probable orthologs for further analysis. More broadly, the power to do this alignment against all prior prokaryotic sequences will enable a wide range of specific analyses, just as BLAST has done with smaller datasets. Some concrete examples were seen in how the approximate (*k*-mer based) matching to the 661k dataset^14^ has been used to search for plasmids^15–19^, adhesins^19^, diversity of vaccine targets^20^, mutations of interest^21^, and phages^22^.

This use-case differs from mapping-to-one-reference in two regards. First, the scale of the intended database in terms of number and diversity. For example, the GTDB r214 representative set^23^, one genome for each of ∼85 thousand bacterial species, contains 242 billion unique *k*-mers - about 65 thousand times more than in one genome. The diversity of gene content of bacteria is very large because many have “open” pangenomes^24,25^, so every new genome adds novel sequence content. Other natural databases to query are larger, but heavily oversample pathogen species - combined GenBank^26^ and RefSeq^27^ contain 2.3 million genomes, and AllTheBacteria^28^ contains 1.8 million high-quality genomes. Thus, there is a computational challenge to index these large databases. Secondly, if mapping to one reference, one try to find the single most likely source of a query, but rarely report all alignments; by contrast, in the BLAST-search use-case, a sequence could truly come from multiple genomes, and all alignments would potentially be wanted by the user.

There are a number of high-performance tools which are good options for large-scale alignment. MMseqs2^29^ supports sensitive and scalable search of nucleotide sequences by searching translated nucleotide databases using a translated nucleotide query. Minimap2^4^, a long-read alignment tool mainly designed for single large reference genome, can also be used for alignment against large scale of microbial genomes, as it can partition input sequences and sequentially index and search each partition. Two tools have demonstrated ability to scale to huge databases, each with a caveat we would want to transcend. First, Phylign^13^ compresses genomes by leveraging phylogenetic information and then given a query, employs a *k*-mer based method COBS^30^ to prefilter genomes before performing base-level alignment with Minimap2. However, as we will show below, prefiltering based on matching 31-mers is only effective for highly similar sequences; as the divergence of the query increases beyond 10%, it becomes very likely to fail the prefilter^31^. It is therefore essential for most useful searches to avoid this limitation. Second, it was shown that by restricting to the 179/11264 species in AllTheBacteria^28^ that have >200 genomes (which is 94% of the data), a new BWT implementation, Ropebwt3^32^, can leverage the within-species redundancy and compress the data from ∼2.8 TB (gzip-compressed) to just 27.6 GB. However, for our use case it is important to be able to align to all the species in the dataset.

In this study, we look at the problem from a fresh perspective. In order to be able to do alignment, we need to be able to find anchoring matches shared between a query and a genome, between which we can do straightforward alignment. We create a relatively small set of probes (20,000 *k*-mers, much smaller than the 59 billion *k*-mers in AllTheBacteria, or 292 billion in GTDB complete dataset) that “cover” the genomes in the database in the sense that every 250 bp window contains several (median 5) *k*-mers, each with a 7 bp prefix-match with one of our probes. Combining this core idea with a range of computational innovations, detailed below, we are able to develop a standalone alignment tool, LexicMap, which is able to align a gene to millions of genomes in minutes.

## Results

### Accurate seeding algorithm

In LexicMap, we first reimplement the sequence sketching method LexicHash^33^, which supports variable-length substring matches (prefix matching) rather than fixed-length *k*-mers, and use this to compute alignment seeds. We outline the approach here and give full details in the Methods. First, twenty thousand 31-mers (called “probes”) are generated (Fig.1a), which can “capture” any DNA sequence via prefix matching, as the probes contain all possible 7-bp prefixes. Then, for every reference genome, each probe captures one *k*-mer across the genome as a seed – this is done via the LexicHash, which chooses the *k*-mer that shares the longest prefix with the probe (Fig.1b, Fig.S2). The number 20,000 above was chosen as a trade-off between index size, alignment accuracy and performance (details below, and Table.S1, Table.S2, and Fig.S1).

**Fig. 1.**
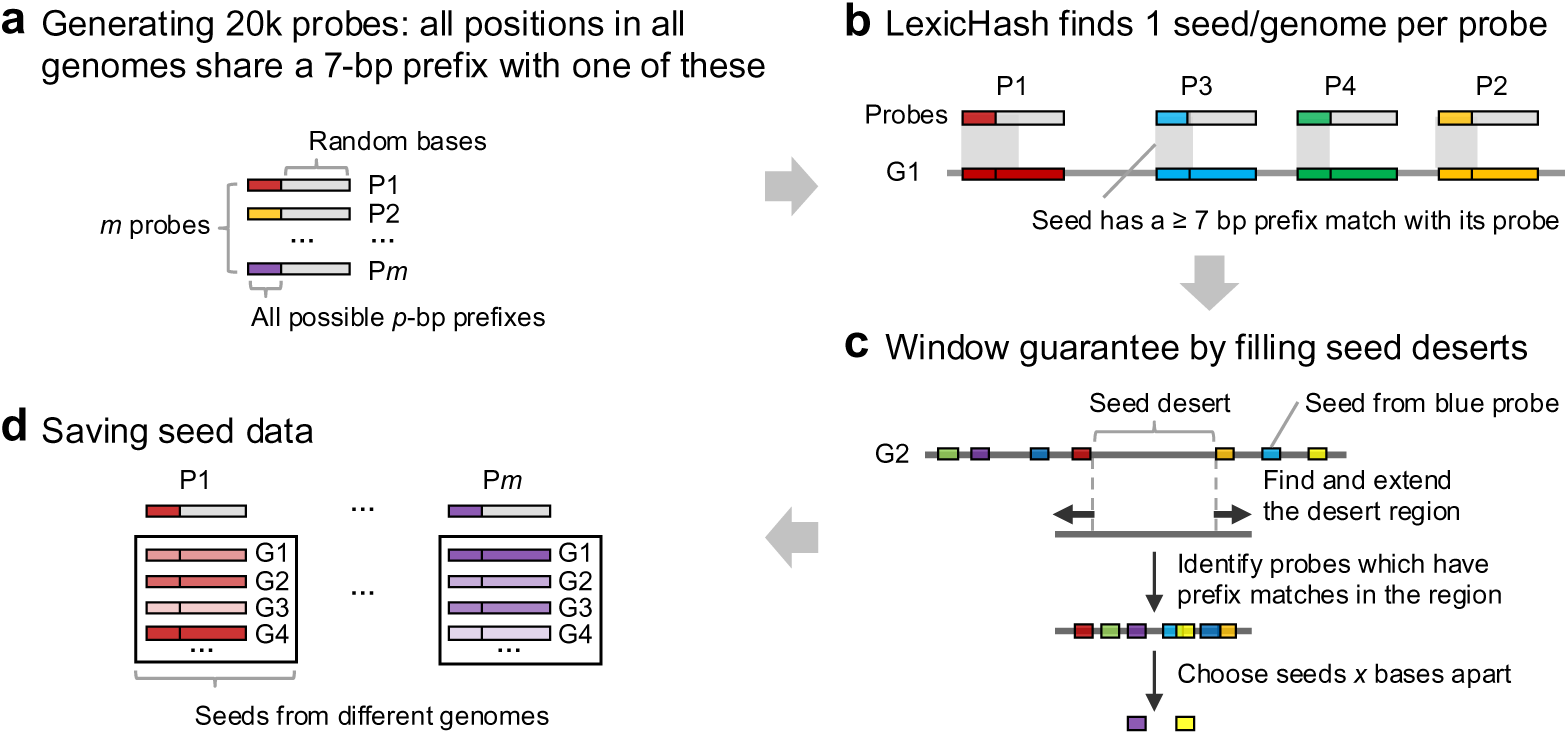
Seeding scheme of LexicMap for reference database. **a)** A fixed set of 20,000 31-mers (called “probes”) are generated, ensuring that their prefixes include every possible 7-mer. Seeds, each prefix-matching one of these, will be found distributed across all database genomes, and chosen in such a way as to have a window guarantee. **b)** LexicHash creates one hash function per probe and when applied to a genome, it finds the *k*-mer with the longest prefix match, which is then stored as a seed. **c)** Each genome is scanned to find seed deserts (regions longer than 100 bp with no seed); every *k*-mer within this region has a 7-mer prefix match with at least one probe (because the probes cover all possible 7-mers), and so seeds can be chosen with spacing of *x* bp (50 by default). **d)** Seeds are stored in a hierarchical index. In fact, although not shown here for simplicity, the number of seeds is doubled to support both prefix and suffix matching (details in Methods).

However, since the captured *k*-mers (seeds) are randomly distributed across a genome, the distances between seeds vary, and there is initially no distance guarantee for successive seeds (Fig.S3a). As a result, some genome regions might not be covered with any seed, creating “seed deserts” or sketching deserts^34^, where sequences homologous to these regions could fail to align. Generally, seed deserts sizes and numbers increase with larger genome sizes (Fig.S3a). To address this issue, a second round of seed capture is performed for each seed desert region longer than 100 bp. New seeds spaced about 50 bp apart are added to the seed list of the corresponding probe (Fig.1c and Fig.1d). After filling these seed deserts, a seed distance of 100 bp is guaranteed (Fig.S3b) for non-low-complexity regions, ensuring that all 250-bp sliding windows contain a minimum of 2 seeds, with a median of 5 in practice (Fig.S3c). Additionally, the seed number of a genome may in practice be lower or higher than the number of probes; the seed number is generally linearly correlated with genome size (Fig.S4). Finally, by allowing variable length prefix-matches, we significantly increase sensitivity compared with a *k*-mer-exact-matching approach, but the method remains vulnerable to variation within the prefix region; we therefore extended LexicMap to additionally support suffix-matching (see Methods).

### Scalable indexing strategies

To scale to millions of prokaryotic genomes, input genomes are indexed in batches to limit memory consumption, with all batches merged at the end (Fig.S5a). Within each batch, multiple sequences (contigs or scaffolds) of a genome are concatenated with 1-kb intervals of N’s to reduce the sequence scale for indexing. Original coordinates and sequence identifiers are restored after sequence alignment. The complete genome or the concatenated contigs are then used to compute seeds using the generated probes as described above, with intervals and gap regions skipped. Genome sequences are saved in a bit-packed format along with genome information for fast random subsequence extraction. After indexing all reference genomes, each probe captures up to millions of *k*-mers, including position information (genome ID, coordinate, and strand). A scalable hierarchical index compresses and stores seed data for all probes (see Methods and Fig.S5a) and supports fast, low-memory variable-length seed matching, including both prefix and suffix matching (Methods and Fig.S5b).

### Efficient variable-length seed matching and alignment

In the searching step, probes from the LexicMap index are used to capture *k*-mers from the query sequence (Fig.2a). Each captured *k*-mer is then searched in the seed data of the corresponding probe to identify seeds that share prefixes or suffixes of at least 15 bp (chosen as a tradeoff between alignment accuracy and efficiency, see Table.S3 and Fig.S6), using a fast and low-memory approach (Methods, Fig.2b). The common prefix or suffix of the query and target seed, along with the position information, constitutes an anchor. These anchors are grouped by genome ID before chaining. The minimum anchor length of 15 bp ensures search sensitivity, while longer anchors (up to 31 bp) provide higher specificity. Unlike minimizer-based methods such as Minimap2, which use fixed-length anchors with a small window guarantee between anchors, LexicMap employs variable-length anchors and does not guarantee a fixed window size between anchors. Consequently, the chaining function (Function 1 in Methods section Chaining) assigns more weight to longer anchors and does not consider anchor distance (see Methods). Next, a pseudo-alignment is performed to identify similar regions from the extended chained regions (Fig.2d). Finally, the Wavefront alignment algorithm is used for base-level alignment (Fig.2e). LexicMap’s default output is a tab-delimited table providing alignment details (Table.S4) for filtration or further analysis, and it also supports an intuitive Blast-style pairwise alignment format (Fig.S7).

**Fig. 2.**
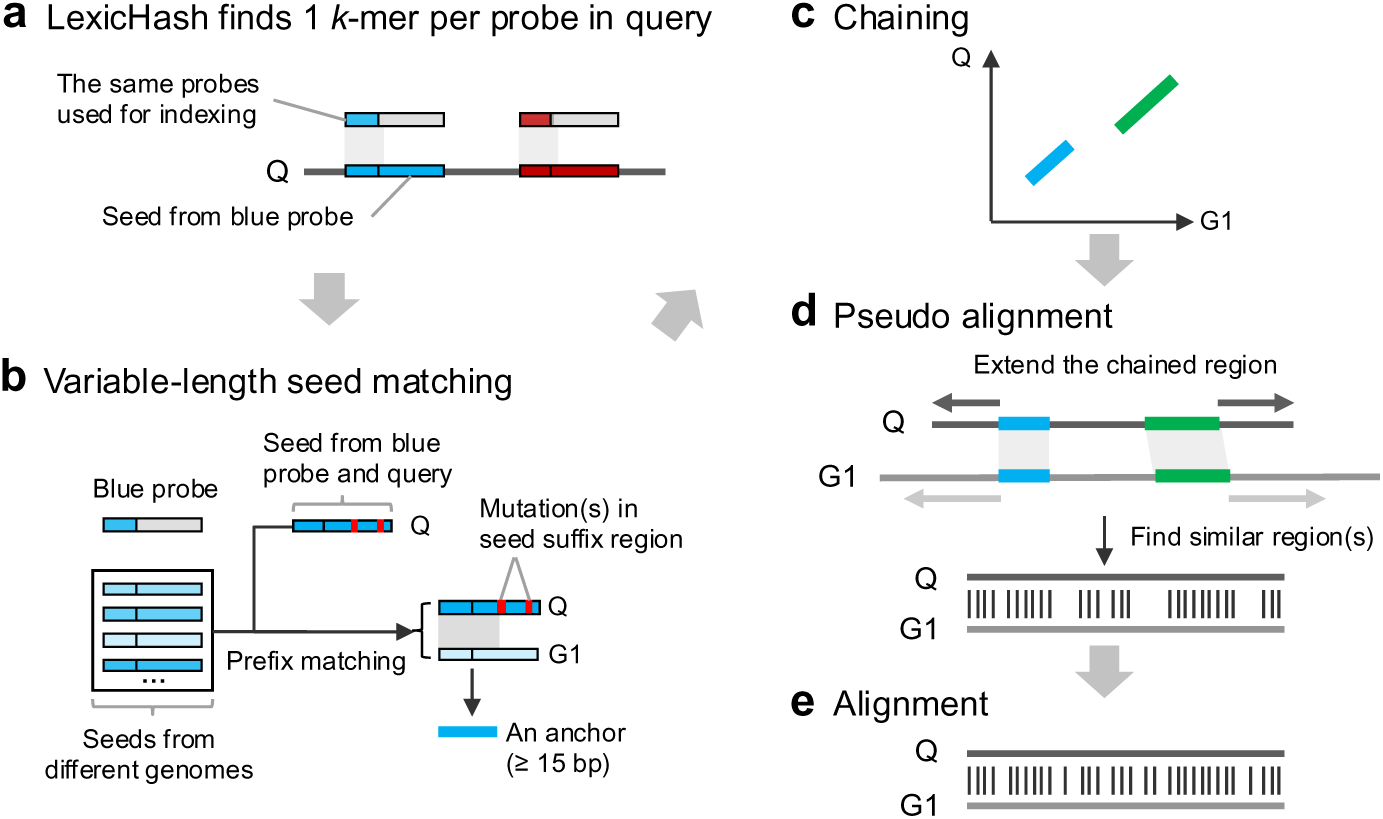
LexicMap alignment workflow. **a)** The same LexicHash hash functions (one per probe) used in the indexing step, are used here, applied to the query to capture one prefix-matching *k*-mer per probe. **b)** For each probe, scan the seed data for that probe, to find prefix or suffix matches ≥ 15bp. The common prefix or suffix constitutes an anchor. **c)** Chain the variable-length anchors using a modified version of the Minimap2 algorithm. **d)** Fast pseudoalignment is followed by **e)** base-level alignment using the Wavefront alignment algorithm. Note that in (b), only prefix matching is illustrated, and suffix matching is not shown for simplicity.

### Robustness to sequence divergence

LexicMap supports variable-length seed matches through prefix and suffix matching, allowing greater tolerance to mutations compared to fixed-length seeding methods. To evaluate LexicMap’s robustness to sequence divergence, ten bacterial genomes from common species with sizes ranging from 2.1 to 6.3 Mb (more details in Table.S5) were used to simulate queries of varying lengths and similarities by introducing SNPs and indels with Badread^35^ (see Methods). Blastn (with word sizes of both the default 28 and 15), MMseqs2, Minimap2, Ropebwt3, were compared with LexicMap. Additionally, COBS was compared as it is used in the prefilter step of Phylign, with a high sensitivity setting (minimum fraction of aligned *k*-mers: 0.33).

Generally, as query identity increases, alignment rates of all tools improve, reaching nearly 100% for query identities ≥ 95% when query length is ≥ 500 bp (Fig.3 and Table.S7). For queries of 1000 bp and 2000 bp, Blastn with a word size of 15 bp (abbreviated as Blastn (ws=15)) consistently achieves the highest alignment rates at lower query identities, followed by MMseqs2, Ropebwt3, Minimap2, LexicMap, and Blastn with the default word size of 28 bp. COBS shows a steeper drop-off in alignment rates at query identities below 95%, which is expected given that it relies on comparing fractions of matched *k*-mers (*k* = 31). For 250 and 500 bp queries, LexicMap outperforms default Blastn at query identities below 93% and 92%, and surpasses Minimap2 at query identities below 88 and 83% respectively. Performance for mutation-free queries is shown in Fig.S8.

**Fig. 3.**
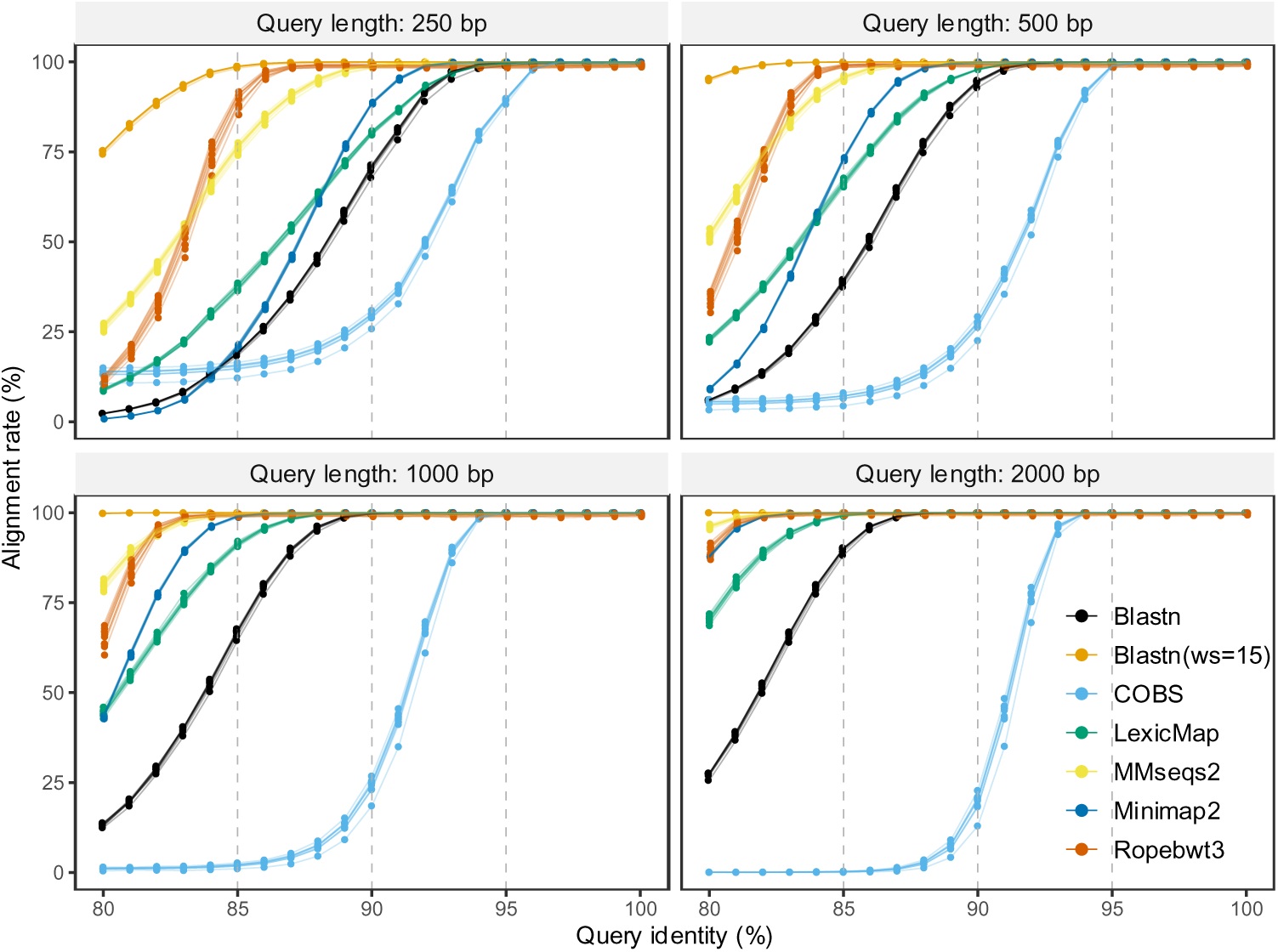
Robustness of aligners to sequence divergence. This is measured by simulating 250, 500, 1000, and 2000 bp reads with coverage of 30× from 10 bacterial genomes, adding mutations to achieve sequence divergence between 0 and 20%, and then aligning back to the source genome. COBS is a *k*-mer index, so here we show what proportion of the “reads” was detected as being present in the source genome (rather than being aligned); this falls off very rapidly as similarity drops because each base disagreement loses an entire 31-mer. For the other tools, we measure the proportion of reads that are correctly aligned back to the source genome. Blastn (ws=15) stands for Blastn with a word size of 15. Blastn (with no brackets) refers to the default setting of Blastn, with word size of 28. All data are available in Table.S7.

### Scalability to 1 million genomes

To evaluate the scalability of the above sequence alignment and search tools to increasing prokaryotic genomes, we created seven genome sets at varying scales (ranging from one to one million genomes) by randomly selecting non-overlapping prokaryotic genomes from GenBank and RefSeq databases. Next, we built an index per set with each tool and then performed searches using a query set containing a rare gene (SecY) and a 16S rRNA gene sequence (see methods).

For index building, generally, the index sizes of all tools are linear correlated with the number of genomes (Fig.4a). In terms of memory requirements to index 1 million genomes, Ropebwt3 requires the most memory (1,013 GB), followed by COBS (382 GB), Minimap2 (85 GB), LexicMap (75 GB), MMseqs2 (20 GB), and Blastn (2 GB). The indexing time of all tools vary (details in Table.S8), ranging from 2.6 hours (MMseqs2) to 23.3 days (Ropebwt3). For databases larger than 10,000 genomes, LexicMap outperforms all other alignment tools in terms of alignment time and memory usage. Note the log scale on the Y axis of Fig.4. For databases of 1 million genomes, LexicMap is 3× faster than the second fast alignment tool (Ropebwt3) while using only 1/115 of the memory (6.2 GB vs 717 GB); it is 89x faster than MMseqs2 and 39x faster than Minimap2.

**Fig. 4.**
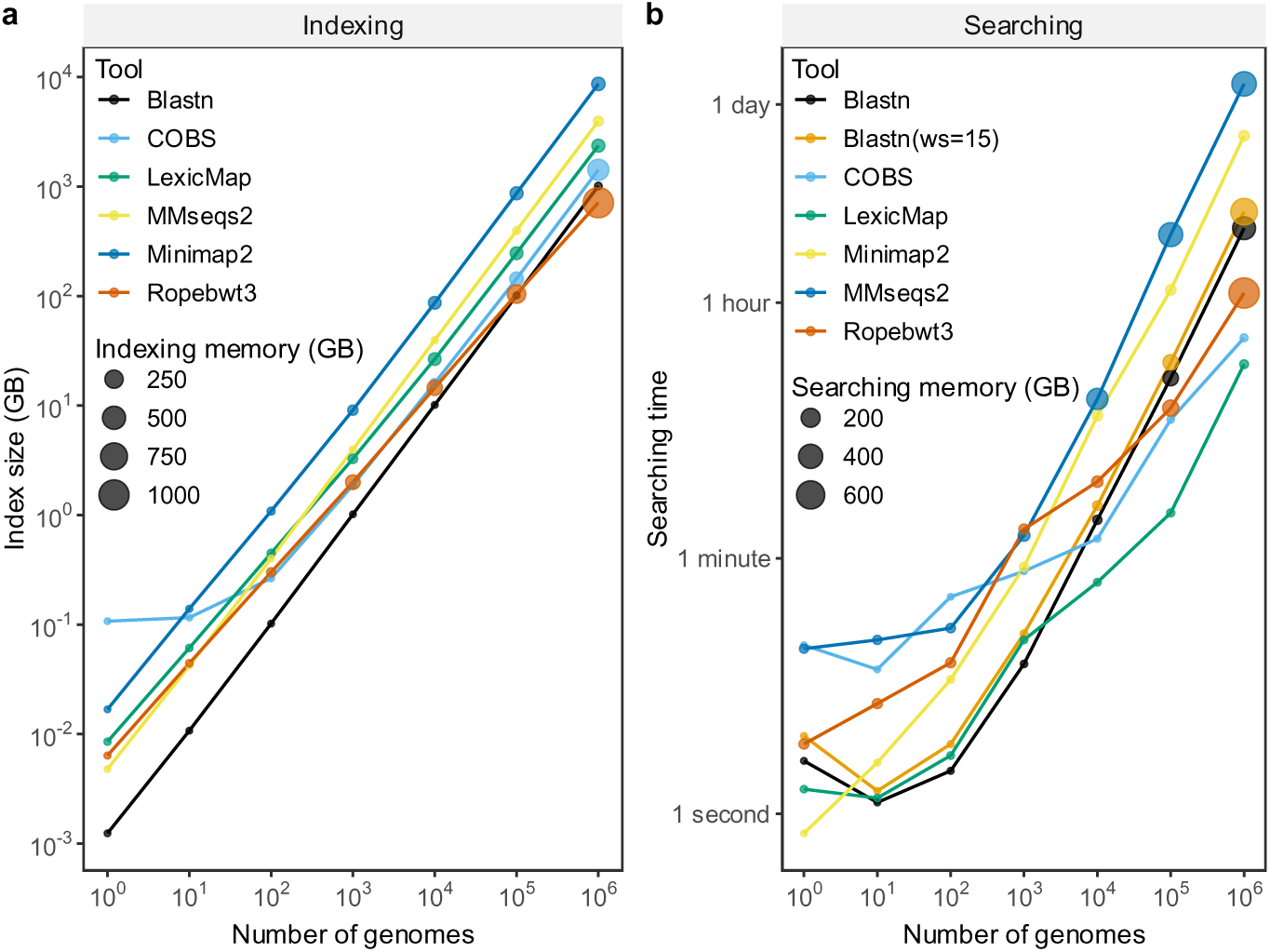
Scalability of sequence alignment/search tools. Benchmarking Blastn, COBS, LexicMap, MMseqs2, Minimap2 and ropebwt3 at construction and then querying of indexes of datasets ranging from 1 genome to 1 million genome**s. a)** Index size and memory requirements for index construction. **b)** Search (meaning alignment for Blastn, LexicMap, MMseqs2, Minimap2, Ropebwt3, and search for COBS) time and memory use. The query set consists of a rare gene (SecY) and a 16S rRNA gene sequence. All tools return all possible matches. Performance data are in Table.S8 and S9.

### Indexing performance on large databases

We further evaluated the performance and accuracy of LexicMap in the three largest and most diverse datasets (see Methods): the GTDB r214 complete dataset with 402,538 prokaryotic assemblies, the AllTheBacteria v0.2 high-quality dataset with 1,858,610 bacterial assemblies, and GenBank+RefSeq with 2,340,672 prokaryotic assemblies (downloaded on February 15, 2024). We benchmarked against Blastn, Minimap2, MMseqs2 and Phylign. Ropebwt3 was excluded from this benchmark because it does not scale to this dataset (see memory use for indexing and querying in the scaling section above) or return all alignments.

In terms of index sizes for AllTheBacteria, Phylign has the smallest size (Table 1), followed by Blastn. LexicMap, MMseqs2, and Minimap2 have index sizes approximately 2.5, 4, and 8 times larger than Blastn, respectively. For indexing time, MMseqs2 is the fastest, followed by Blastn, Minimap2, and LexicMap. For indexing memory, Blastn uses the minimum memory, followed by MMseqs2, Minimap2, and LexicMap. LexicMap uses almost twice as much memory for the GenBank+RefSeq dataset compared to the AllTheBacteria dataset, as the genomes in the former are more diverse.

**Table 1.**
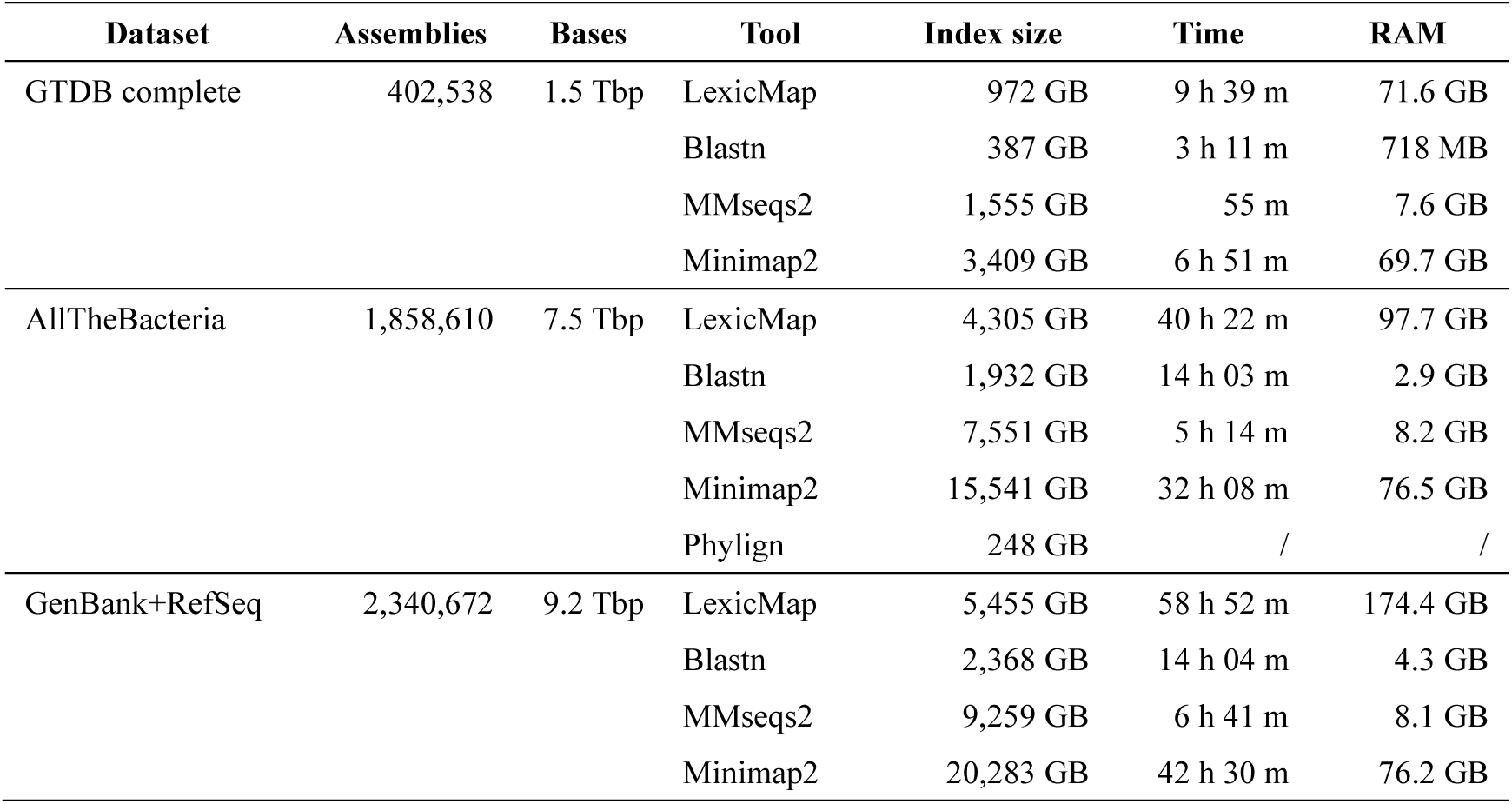
LexicMap indexing performance on three datasets. Tbp: terabase; MB: megabyte (base 1000); GB: gigabyte (base 1000). LexicMap built indexes with the genome batch size of 5,000 for GTDB and 25,000 for the AllTheBacteria dataset and the GenBank+RefSeq dataset.

### Alignment accuracy and performance on large databases

Four different types of queries were used to evaluate alignment performance (see Methods): a) a comparatively rare gene (Blastn returns seven thousand genome hits in GTDB r214 complete dataset with 402,538 genomes) - SecY from *Enterococcus faecalis*, b) a 16S rRNA gene from *Escherichia coli*, c) a 53kb plasmid, and d) 1033 different AMR genes (batch queries).

First, we aligned the queries with LexicMap, Blastn, MMseqs2, and Minimap2 against the GTDB complete index. For clearer comparison, we divided all alignment results into three groups (high, medium and low similarity) according to the query coverage and percentage identities and counted the number of correctly matched genomes (genome hits) – see Table 2.

**Table 2.** Alignment performance benchmarks on GTDB complete dataset. Hits stand for genome hits. Hits (high), hits (medium), and hits (low) mean the number of genomes with high-, medium-, and low-similarity matches, respectively. The details of alignment result grouping are available in Methods.

| Query (length) | Tool | Hits (total) | Hits (high) | Hits (medium) | Hits (low) | Time | RAM |
| --- | --- | --- | --- | --- | --- | --- | --- |
| A rare gene<br>1,299 bp | LexicMap | 6,255 | 2,311 | 46 | 3,898 | 30 s | 2.1 GB |
|  | Blastn | 7,121 | 2,311 | 47 | 4,763 | 2,171 s | 351.2 GB |
|  | Blastn(ws=15) | 57,741 | 2,311 | 47 | 55,383 | 3,171 s | 324.1 GB |
|  | MMseqs2 | 67,537 | 2,304 | 54 | 65,179 | 26,174 s | 400.7 GB |
|  | Minimap2 | 2,312 | 2,312 | 0 | 0 | 17,208 s | 20.2 GB |
| A 16S rRNA gene<br>1,542 bp | LexicMap | 306,064 | 60,999 | 69,293 | 175,772 | 303 s | 5.2 GB |
|  | Blastn | 301,197 | 61,878 | 109,477 | 129,842 | 2,760 s | 378.4 GB |
|  | Blastn(ws=15) | 301,197 | 61,878 | 109,477 | 129,842 | 3,291 s | 378.4 GB |
|  | MMseqs2 | 324,364 | 60,915 | 89,874 | 173,575 | 31,140 s | 400.7 GB |
|  | Minimap2 | 17,656 | 15,998 | 1,652 | 6 | 17,313 s | 20.2 GB |
| A plasmid<br>52,830 bp | LexicMap | 65,029 | 21 | 2,808 | 62,200 | 539 s | 6.8 GB |
|  | Blastn | 69,311 | 21 | 2,865 | 66,425 | 2,262 s | 364.7 GB |
|  | Blastn(ws=15) | 91,847 | 21 | 2,865 | 88,961 | 3,082 s | 142.8 GB |
|  | MMseqs2 | 90,277 | 7 | 1,650 | 88,620 | 44,710 s | 400.7 GB |
|  | Minimap2 | 3,033 | 35 | 1,873 | 1,125 | 19,715 s | 20.2 GB |
| 1033 AMR genes<br>1 kb (median) | LexicMap | 4,665,317 | 1,123,251 | 776,153 | 2,765,913 | 8,620 s | 10.7 GB |
|  | Blastn | 5,357,772 | 1,150,407 | 772,858 | 3,434,507 | 4,686 s | 442.1 GB |
|  | Blastn(ws=15) | 10,877,544 | 1,150,410 | 840,464 | 8,886,670 | 4,561 s | 311.9 GB |
|  | MMseqs2 | 10,137,345 | 1,148,942 | 808,177 | 8,180,226 | 184,470 s | 406.9 GB |
|  | Minimap2 | 2,078,490 | 943,516 | 39,529 | 815,445 | 38,058 s | 20.2 GB |

In short – all tools find very similar numbers of high-similarity alignments, but LexicMap reports fewer low-similarity/fragmentary alignments. For the rare gene all tools return almost identical numbers of high-similarity alignments, but MMseqs2 and Blastn report about 10× more low similarity alignments than other tools. For the 16S rRNA gene, where we expect to find many alignments, LexicMap, Blastn (both settings) and MMseqs2 report ∼61k high-similarity alignments, whereas Minimap2 only report ∼16k. Counting all (high/medium/low-similarity) alignments, again all tools except Minimap2 found around 300k alignments (MMseqs2 found the most, 324k), whereas Minimap2 reports ∼18k. In contrast, the 53kb plasmid, which is likely to present a different type of challenge, with many small fragmentary hits and some long hits with large deletions, revealed other differences between the tools. LexicMap and Blastn (both settings) find 21 high-similarity matches, and Minimap2 finds 35, but MMseqs2 finds only 7. However Blastn (ws=15) and MMseqs2 find many more low-similarity alignments than other tools. Finally for the AMR genes, LexicMap, Blastn (both settings) and MMseqs2 find around 1.1 million alignments, whereas Minimap2 finds 943k. However again Blastn (ws=15) and MMseqs2 report 4× more low-similarity alignments.

In terms of speed, LexicMap is significantly faster than other tools for single queries, being 72×, 9×, and 4× faster than the second fastest tool Blastn, on the rare gene, 16S rRNA gene and plasmid respectively. Compared with MMseqs2, LexicMap is 872×, 103×, 83× faster for the same queries. For batch queries, LexicMap is 1.8× slower than Blastn. Regarding memory usage, LexicMap required less than 7 GB for single queries and 11 GB for 1033 AMR genes, whereas Minimap2 uses 20.2 GB and Blastn and MMseqs2 use more than 300 GB across all queries.

Next, we compared LexicMap with Phylign on the AllTheBacteria dataset, which contains 1,858,610 bacterial genomes, including some species that are highly over-sampled. Here, Blastn could not be run due to its requirement of more than 2000 GB of memory. MMseqs2 was not included for its slow speed, and Minimap2 was not included for its slow speed and lower sensitivity for medium- and low-similarity matches. Across all queries, if including all (high/medium/low-similarity) hits, LexicMap returns more genome hits than Phylign, but for high-similarity matches, the number of alignments is very similar. These observations are as expected given the effect of Phylign using a *k*-mer filter on diverged hits (see Fig.3). In terms of computation efficiency, for single queries, LexicMap takes much less time than Phylign both in local and cluster mode (using up to 100 nodes), while also using much less memory. However for batch querying, LexicMap is much slower than Phylign in local and cluster modes, although in both cases LexicMap returns more alignments.

Finally, we tested LexicMap on GenBank+Refseq, where it achieved similar performance on the GenBank+RefSeq dataset (234 million prokaryotic genomes) as it does on the AllTheBacteria dataset (Table S6).

**Table 3.** Alignment performance benchmarks on AllTheBacteria high-quality dataset*.

| Query (length) | Tool | Hits (total) | Hits (high) | Hits (medi) | Hits (low) | Time | RAM |
| --- | --- | --- | --- | --- | --- | --- | --- |
| A rare gene | LexicMap | 38,062 | 7,935 | 18 | 30,109 | 112 s | 3.8 GB |
| 1,299 bp | Phylign_local | 7,937 | 7,935 | 1 | 1 | 1,848 s | 77.6 GB |
|  | Phylign_cluster | 7,937 | 7,935 | 1 | 1 | 1,713 s | / |
| A 16S rRNA gene | LexicMap | 1,857,974 | 496,867 | 556,951 | 804,156 | 1,552 s | 14.8 GB |
| 1,542 bp | Phylign_local | 1,017,766 | 483,054 | 434,105 | 100,607 | 7,833 s | 77.0 GB |
|  | Phylign_cluster | 1,017,766 | 483,054 | 434,105 | 100,607 | 5,141 s | / |
| A plasmid | LexicMap | 485,295 | 25 | 9,201 | 476,069 | 1,891 s | 15.2 GB |
| 52,830 bp | Phylign_local | 46,822 | 27 | 9,832 | 36,963 | 2,853 s | 82.6 GB |
|  | Phylign_cluster | 46,822 | 27 | 9,832 | 36,963 | 2,374 s | / |
| 1033 AMR genes | LexicMap | 25,563,227 | 6,693,084 | 3,814,828 | 15,055,315 | 44,231 s | 17.7 GB |
| 1 kb (median) | Phylign_local | 11,742,865 | 5,796,412 | 2,871,952 | 3,074,501 | 9,488 s | 85.9 GB |
|  | Phylign_cluster | 11,742,865 | 5,796,412 | 2,871,952 | 3,074,501 | 8,029 s | / |

\* Alignment memory can not be accurately measured for Phylign running in multiple cluster nodes.

## Discussion

BLAST was not the first tool to enable DNA alignment against a database, but its speed and accessibility revolutionised bioinformatics. However since then, NCBI BLAST has been querying an exponentially smaller fraction of public data^13^. The vast majority of the cellular tree of life consists of bacteria and archaea; we continue to expand the tree through metagenomic sequencing, and although the rate of discovery of new phyla has dropped, this is not the case for lower taxa^25^. Taken together with the prevalence of horizontal gene transfer in prokaryotes and a high number of mobile genetic elements and their cargo, the levels of diversity are extremely high and continue to grow. Further, the amount of clinical sequencing and deposition in archives continues to grow our collection of pathogen sequence data, providing a year-by-year perspective on real-time evolution and horizontal gene transfer. Thus now more than ever, the ability to align a query against a database of representative genomes, or all genomes, is vital to modern biology and public health. Developers of new antibiotics who find specific mutations confer resistance should be able to find out whether those SNPs have been seen before, representing pre-existing resistance^36^. Genomic epidemiologists should be able to query a drug-resistance plasmid from a hospital outbreak against recently sequenced genomes from across the world^15^ or track down any global samples containing outbreak informative SNPs^37^. Just as BLAST enabled hundreds of different studies that could not have been predicted at the time, re-enabling alignment against all bacteria will surely potentiate a wide range of new applications.

We have introduced our solution to this problem here. LexicMap takes a novel approach, constructing a fixed set of 20,000 probes (*k*-mers) which are guaranteed to have multiple (around 5 in this study) prefix matches in every 250 bp window of every genome in the database. The *k*-mer with the best prefix match for each probe in each genome is (called a seed and) stored in a hierarchical index, which can be used for alignment. This use of variable-length prefix-and suffix-matching enables sensitive nucleotide alignment for queries above 250 bp long (though this is a user-tunable threshold). Our results show (Fig. 4) that LexicMap achieves a superior scalability to the other benchmarked alignment tools, with the fastest speed and the lowest memory usage, while maintaining a moderate index size and indexing efficiency. While achieving this scalability, LexicMap maintains a comparable sensitivity (meaning robustness to divergence of the query from the target) to state-of-the-art aligners.

In terms of user experience, the main advantages of LexicMap compared with current approaches include: 1) It provides direct alignment without a lossy prefilter step. 2) All possible matches, including multiple copies of genes in a genome, are returned. 3) The alignment is fast and memory efficient. 4) Unlike minimizer-based methods, seeds in LexicMap are interpretable and several utility commands are available to interpret probe (probe *k*-mers) and seed data (seed sequences and positions). 5) LexicMap is easy to install on multiple operating systems and can be used as a standalone tool without needing a workflow manager or compute cluster. LexicMap mainly supports small genomes including complete or partially assembled prokaryotic, viral, and fungi genomes. The maximum supported sequence length is 268,435,456 bp (2^28^), so it can in principle also be applied to bigger genomes like human genomes with a maximum chromosome size of 248 Mb.

In terms of limitations, LexicMap only supports queries longer than 250 bp. It achieves a low memory footprint by storing a very large index on disk (5.46 TB for 2.34 million prokaryotic assemblies in GenBank+RefSeq), although we do note that the corresponding index for MMseqs2 or Minimap2 would be larger. Nevertheless, it would be desirable in future to reduce the size of this index. Finally, LexicMap is optimized for a small number of queries; improving batch searching speed is planned in the future.

Large-scale alignment against representative and complete genome databases has been a vital part of biology for decades, and as sequencing has accelerated, the need for scalable solutions has grown. LexicMap marks a step-change in scalability, achieving low-memory queries of the global corpus of bacterial data in minutes. Coupled with its ease-of-installation and use, without the need for workflow managers or a cluster, it has the potential to enable a wide range of analyses from ecology, evolution and epidemiology.

## Online Methods

### Probe generation

Probes, also referred to as “masks” in the LexicHash paper, consist of a fixed number (*m* = 20,000 by default) of *k*-mers (*k* ≤ 32, *k* = 31 by default), which “capture” DNA sequences via prefix matching. A probe consists of two parts – the *p*-bp prefix and the remaining bases. To enable probes to match all possible reference and query sequences via prefix matching, all permutations of *p*-mers are generated as the base prefix set. The length of the prefix (*p*) is calculated to ensure that the total number of possible prefixes (4^p^) does not exceed the total number of probes (*m*). For instance, when *m* = 20,000, *p* equals 7. Then, the base prefixes are duplicated to reach the number of probes *m*. Next, the suffixes of probes are randomly generated. The *p’*-bp (*p’* = *p* + 1) prefixes are required to be distinct to enable fast locating of the probe via the prefix of a *k*-mer through an array data structure. Finally, these *m k*-mers serve as probes that can match all potential sequence regions in both reference genomes and query sequences.

### Seed computation

*K*-mers are encoded as 64-bit unsigned integers, with a binary coding scheme of A=00, C=01, G=10, and T=11. Following the original implementation of LexicHash, the hash value between a probe and a *k*-mer is computed using a bitwise XOR operation. The value of this hash is that smaller hash values indicate longer common prefixes between the probe and the *k*-mer, while a hash value of zero means the probe and the *k*-mer are identical.

For each reference genome or query sequence, all *k*-mers from both forward and backward strands are compared with probes that share the same *p*-bp prefix. Each probe retains only the *k*-mer with the minimum hash value, which might in principle be located at more than one position. *K*-mers of low-complexity are discarded by DUST algorithm^38^ with a loose score threshold of 50. A 64-bit integer encodes each position’s information, including the genome batch index (17 bits, see “indexing” section), genome index (17 bits), position (28 bits), the strand (1 bit), and the seed direction flag (1 bit, see “Seed data storage and variable-length seed matching” section). Ultimately, each probe captures one *k*-mer (as a seed) across the entire reference genome or query sequence, which shares the longest prefix with the probe.

However, due to the random distribution of captured *k*-mers (seeds), the distances between these seeds vary, and there is no guarantee of consistent distance between successive seeds. Consequently, some regions of the sequence might remain uncovered by seeds, leading to what are known as “seed deserts”. These deserts are problematic because they can cause sequences homologous to the regions to fail to align. To address this issue, regions identified as seed deserts, which are longer than a certain threshold (100 bp by default), are extended by 1 kb both upstream and downstream. A second round of seed capture is then performed in these extended regions, and new seeds spaced about *x* (50 by default) bp apart within the region are added to the index of the corresponding probes. After filling these seed deserts, the total number of seeds may exceed the initial value of *m*.

### Indexing

The input of LexicMap is a list of microbial genomes, with the sequences of each genome stored in separate FASTA files. These files can be in plain or compressed formats such as gzip, xz, zstd, or bzip2. Each file must have a distinct genome identifier in the file name. To limit memory consumption, genomes are indexed in batches, and all batches are merged at the end (Fig.S5a).

In each batch, genomes with any sequence larger than a genome size threshold (15 Mb by default), such as non-isolate assemblies, are skipped. While if only the total length of sequences exceeds the threshold, the genomes are split into multiple chunks and alignments from these chunks will be merged in the searching step. Additionally, unwanted sequences within genomes, such as plasmids, can be optionally discarded using regular expressions to match sequence names. Multiple sequences (contigs or scaffolds) of a genome are concatenated with 1-kb intervals of N’s, to reduce the scale of sequences to be indexed. The original coordinates will be restored after sequence alignment. The complete genome or the concatenated contigs are then used to compute seeds with generated probes, as described above. Before this, any degenerated bases are converted to their corresponding alphabet-first bases (e.g., N is converted to A). The genome sequence is then saved in a bit-packed format (2 bits per base), along with associated genome information (genome ID, size, and sequence IDs and lengths of all contigs) to enable fast random subsequence extraction. Simultaneously, seeds and their positional information are appended to their corresponding probes and saved as seed files. After processing all genomes in the batch, these seed files are merged using an external sorting method once all batches have been indexed.

### Seed data storage and variable-length seed matching

After indexing with all *n* reference genomes, each probe captured *n* or more seeds (*k*-mers, encoded as 64-bit integers), with each seed potentially having one or more positions (also represented as 64-bit integers), and there are *m* probes in total (20,000 by default). To scale to millions of prokaryotic genomes, the storage of seed data needs to be both compact and efficient for querying. Since the seeds of different probes are independent, the seed data are saved into *c* chunk files to enable parallel querying (Fig.S5a). In each chunk file, the seed data of approximately *m*/*c* probes are simply concatenated. For each probe, all seeds are sorted in alphabetical order, and the varint-GB^39^ algorithm is used to compress every two seeds along with their associated position counts. Since seeds captured by the same probe share common prefixes, the differences between two successive seeds are small, as are the position counts. As a result, these values can be stored using as little as two bytes instead of 16 bytes.

Unlike the approach in the LexicHash paper, where captured *k*-mers from each probe are stored in a prefix tree in main memory, here they are alphabetically sorted and saved in a list-like structure within files. To enable efficient variable-length prefix matching of seeds, an index is created for each seed data file (Fig.S5b). This index functions similarly to a table of contents in a dictionary, storing a list of marker *k*-mers along with their offsets (pages) in the seed data file. Each marker *k*-mer is the first one with a specific *p’’*-bp subsequence (*p’’* = 6 by default) following the *p*-bp prefix (all seeds of a probe share the same *p*-bp prefix). For a query sequence, one *k*-mer is captured by each probe and searched within the corresponding probe’s seed data to return seeds that share a minimum length of prefix with the query *k*-mer. For example, searching with CATGCT for seeds (with *p* = 2, *p’’* = 1) that have at least 4 bp common prefixes is equivalent to finding seeds in the range of <u>CATG</u>AA to <u>CATG</u>TT. The process starts by extracting the *p’’*-bp subsequence from <u>CATG</u>AA (in this case, T) to locate the marker *k*-mer (e.g., CA<u>T</u>CAC) with the same *p’’*-bp subsequence in the same region. The offset information (page) of this marker *k*-mer is then used as the starting point for scanning seeds within the *k*-mer range.

The index structure described above is extended to support the suffix matching of seeds. During the indexing phase, after a seed *k*-mer is saved into the seed data of its corresponding probe, the *k*-mer is reversed and added to the seed data of the probe that shares the longest prefix with the reversed *k*-mer. Additionally, the last bit of each position data is used as a seed direction flag, indicating that the seed *k*-mer is reversed. As a result, all seeds are doubled: there are “forward seeds” for prefix matching and “reversed seeds” for suffix matching, which is achieved through prefix matching of the reversed seeds. In the seed matching process, two rounds of matching are performed. The first round involves prefix matching (as described above) in the forward seed data. In the second round, the query *k*-mer is reversed and searched in the reversed seed data of the probe that shares the longest prefix with the reversed *k*-mer.

### Chaining

After searching all captured *k*-mers from a query in the seed data of all probes, seed pairs (anchors) with different matched prefix/suffix lengths *l* are returned. These anchors indicate matched regions between the query sequences [*y*, *y* + *l* - 1] on the strand *s*_*y*_ and reference genome [*x*, *x* + *l* - 1] on strand *s*_*x*_. First, the anchors are grouped by the genome IDs. Within each group, the anchors are sorted by the following criteria: 1) Ascending order of start positions in the query sequence. 2) Descending order of end positions in the query sequence. 3) Ascending order of start positions in the target genome. Only one anchor is kept for duplicated anchors, and any inner anchors that are nested within other anchors are removed. Then, overlapped anchors with no gaps are merged to a longer one, while for these with gaps, only the non-overlapped part of the second anchor is used to compute the weight (see below). Next, a chaining function modified from Minimap2 is applied to chain all possible colinear anchors:

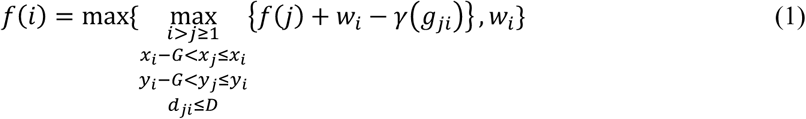

Here, *f*(*i*), *f*(*j*) are scores for anchors *a*_*i*_ and *a*_*j*_, respectively. The anchor weight 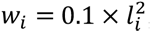, where *l*_*i*_ is the length of the anchor *a*_*i*_ and *P* ≤ *l*_*i*_ ≤ *k*, with *P* being the minimum anchor length (15 by default). The seed distance *d*_*ji*_ = max{|*y*_*i*_ – *y*_*j*_|, |*x*_*i*_ – *x*_*j*_|} and *D* is the maximum seed distance (1,000 by default). The seed gap 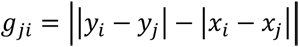, and *G* is the maximum gap (50 by default). The gap penalty is calculated as *γ*(*g*) = 0.1 × *g* + 0.5 log_2_(*g*). Additionally, a filter is applied to avoid non-monotonic increase or decrease in coordinates. For cases where there is only one anchor, the length of the anchor must be no less than a threshold *P’* (17 by default).

### Alignment

In contrast to the case of minimizers or closed syncmers^40^, seeds in LexicMap do not have a small window guarantee. Hence a pseudo alignment algorithm is further used to find similar regions from the chained region, extended by 1-kb in the upstream and downstream directions. *K*-mers (*k* = 31) on both strands of the query sequence are stored in a prefix tree, and *k*-mers of a target region are queried in the prefix tree to return matches of *l*_*i*_^′^ (11 ≤ *l*_*i*_^′^ ≤ 31) in the extended chained region. After removing duplicated anchors as above, the anchors are chained with the score function:

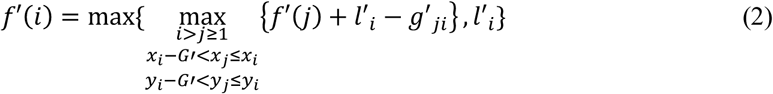

where *G*′ is the maximum gap (20 by default), seed distance 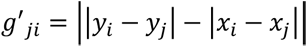. And the chaining is banded (100 bp or 50 anchors by default). Besides, chained regions across the interval regions between contigs are split. Next, regions are further extended in both ends, via a similar pseudo alignment algorithm based on 2-mer matches.

Similar regions between the reference and query sequences are further aligned with a reimplemented Wavefront alignment algorithm (https://github.com/shenwei356/wfa, v0.4.0), with a gap-affine penalty (match: 0, mismatch: 4, gap opening: 6, gap extension: 2) and the adaptive reduction heuristic (minimum wavefront length: 10, maximum distance: 50). Each alignment’s bit score and expect value (e-value) are computed with Karlin-Altschul parameter values for substitution score 2 and −3 (gap opening cost: 5, gap extension cost: 2, lambda: 0.625, K: 0.41) from Blastn’s source code.

### Tools, reference genome datasets and query sequences

LexicMap v0.7.0, Blast++ v2.15.0, MMseqs2 v16.747c6, Minimap2 v2.28-r1209, Ropebwt3 v3.9-r259, COBS (iqbal-lab-org fork v0.3.0, https://github.com/iqbal-lab-org/cobs), and Phylign (AllTheBacteria fork https://github.com/AllTheBacteria/Phylign, commit 9fc65e6) were used for testing. GTDB r214 complete dataset with 402,538 prokaryotic assemblies was downloaded with genome_updater (https://github.com/pirovc/genome_updater, v0.6.3). GenBank+RefSeq dataset with 2,340,672 prokaryotic assemblies was downloaded with genome_updater on February 15, 2024. AllTheBacteria v0.2 high-quality genome dataset with 1,858,610 bacteria genomes and the Phylign index were downloaded from https://github.com/AllTheBacteria/AllTheBacteria. Four query datasets were used for evaluating alignment accuracy and performance: a preprotein translocase subunit SecY gene CDS sequence from an *Enterococcus faecalis* strain (NZ_CP064374.1_cds_WP_002359350.1_906), a 16S rRNA gene from *Escherichia coli* str. K-12 substr. MG1655 (NC_000913.3:4166659-4168200), a plasmid from a *Serratia nevei* strain (CP115019.1), and 1033 AMR genes from the Bacterial Antimicrobial Resistance Reference Gene Database (PRJNA313047). All files were stored on a network-attached storage server equipped with HDD disks.

### Simulating mutations

Ten bacterial genomes of common species (more details in Table.S5) with genome sizes ranging from 2.1∼6.3 Mb were used for simulating queries with Badread^35^ v0.4.1 and SeqKit^41^ v2.8.2. The command, “badread simulate --seed 1 --reference $ref --quantity 30x --length $qlen,0 –identity $ident,$ident,0 --error_model random --qscore_model ideal --glitches 0,0,0 --junk_reads 0 -- random_reads 0 --chimeras 0 --start_adapter_seq ‘’ --end_adapter_seq ‘’ | seqkit seq -g -m $qlen | seqkit grep -i -s -v -p NNNNNNNNNNNNNNNNNNNN -o $ref.i$ident.q$qlen.fastq.gz”, simulated reads with a genome coverage of 30×, a mean length of 250, 500, 1,000 or 2,000 bp and mean percentage identities ranging from 80% to 100% with SNPs and indels. LexicMap, Blastn, MMseqs2, Minimap2, Ropebwt3, and COBS were used to search simulated reads against the reference genomes. Each tool built an index for each genome and searched or aligned reads with the corresponding index. LexicMap built indexes and aligned with default parameters (*m* = 20,000, *k* = 31, *P* = 15, *P’* = 17). Blast built indexes with default parameters and aligned reads with the default task mode “megablast” using a word size of 28 (default) or 15, and the out format 6 was set to output tabular results. MMseqs2 built indexes and aligned reads with the nucleotide mode, and the format mode 4 was set to output tabular results. Minimap2 aligned reads with the “map-ont” mode. Ropebwt3 built indexes (including .fmd, .ssa and .fmd.len.gz files) with default parameters and aligned with the local alignment mode (“ropebwt3 sw”). COBS built compact indexes with *k* = 31 and a false positive rate of 0.3 and searched reads with a *k*-mer coverage threshold of 0.33. Positions of alignment results from all tools except for COBS, were checked, and only those overlapping with the source locations were counted for the computation of alignment rate. Unless otherwise specified, all tests were performed in cluster nodes running with Intel(R) Xeon(R) Gold 6336Y CPU @ 2.40GHz with RAM of 500 or 2000 GB.

### Scalability testing

Seven genome sets with 1, 10, 100, 1,000, 10,000, 100,000, and 1,000,000 non-overlapping prokaryotic genomes randomly choosing from GenBank+RefSeq dataset were used for the scalability test. The queries consist of a SecY and a 16S rRNA gene sequences mentioned above. LexicMap, Blastn, MMseqs2, Minimap2, Ropebwt3, and COBS built an index for each genome set and searched or aligned the queries against the corresponding index, with the options in the previous tests. LexicMap returned all possible matches by default, while other tools were explicitly set to return all matches. All tools used 48 CPUs. A Python script (https://github.com/shenwei356/memusg) was used to record the time and peak memory usage. Sequence alignment/searching was repeated four times on different cluster nodes over separate weeks, and the average time and memory consumption were used for plotting.

### Benchmarking

For indexing, LexicMap built indexes with the genome batch size 5,000 (default) for GTDB and 25,000 for the AllTheBacteria dataset. Blastn, MMseqs2, and Minimap2 build indexes with parameters as mentioned above. The Phylign index was built with default parameters, including *k* = 31 and a false-positive rate of 0.3 for the COBS index; since the index building involved three workflows with multiple steps in multiple cluster nodes, the memory and time could not be measured accurately.

The four query datasets were used for sequence alignment. LexicMap returned all possible matches by default and other tools were set to return all possible matches according to the sequence number in a database. All tools used 48 CPUs. The main parameters of Phylign included threads=48, cobs_kmer_thres=0.33, minimap_preset="asm20", nb_best_hits=5,000,000, max_ram_gb=100. For the cluster mode, the maximum number of Slurm jobs was set to 100.

The four tools’ sequence alignment results were divided into three categories according to query coverage (qcov) and percentage of identity (pident), and the genome number of each category were counted for comparison. The metrics of Minimap2 and Phylign were computed by sam2tsv.py (https://gist.github.com/apcamargo/2b7ca3032c1e80333adc1e54f47a0966). Alignments of high similarity are those with qcov≥90% (genes) or 70% (plasmids) and pident≥90%. Alignments of low similarity are those with qcov<50% (genes) or 30% (plasmid) or pident < 80%. Left alignments are marked as medium similarity.

## Code availability

LexicMap is an open-source standalone tool implemented in Go under the MIT license at https://github.com/shenwei356/LexicMap, with freely available statically linked executable binary files for common operating systems and CPU types. The source code is also archived at https://doi.org/10.5281/zenodo.15197523. Two main subcommands *index* and *search* are used to create an index and perform searching, respectively, and several utility subcommands are available for interpreting the index data and extracting indexed sequences.

## Data availability

All analyses were done with data public genome databases; specifically GTDB r214 complete dataset, GenBank+RefSeq dataset, downloaded on February 15, 2024, and AllTheBacteria v0.2. Full details on how to reproduce all analyses, along with lists of accessions used, can be found in this GitHub repository https://github.com/shenwei356/lexicmap-benchmark, and also in this Zenodo archive: https://doi.org/10.5281/zenodo.15628530.

## Acknowledgements

This study was supported by grants from the National Natural Science Foundation of China (82341112), Chinese Scholarship Council scholarship (202308500105), EMBL Visitor/Sabbatical Programme fellowship, Remarkable Innovation-Clinical Research Project, Joint Project of Pinnacle Disciplinary Group, and Kuanren Talents Program of The Second Affiliated Hospital of Chongqing Medical University.

We thank Shuyi Wang (Peking University People’s Hospital, China), Leah Roberts (Queensland University of Technology, Australia), Songbai Cai and Liuyang Zhao (Chongqing Medical University, China), and Rachel Colquhoun (Edinburgh University, UK) for using LexicMap and giving valuable feedback during the development. We thank Daniel Anderson for suggesting test datasets. We thank Pengfei Wang (University of Montpellier, France) for comments on the manuscript and visualization. We thank Daniel Anderson, Martin Hunt, and Daria Frolova for fruitful discussions.

## Author contributions

W.S. and Z.I. designed the project. Z.I. managed the project. W.S. implemented the software. Z.I. and J.L. provided computing resources. W.S. and Z.I. performed benchmarks and data interpretation. W.S., Z.I., and J.L. wrote the manuscript. All authors reviewed and approved the manuscript.

## Extended Data

**Table. S1.** Index sizes and indexing performances with different numbers of probes. The reference genomes were 402,538 prokaryotic assemblies from GTDB r214 complete dataset. LexicMap `index` command used default parameters except for the number of probes. The tests were performed on a server with Intel(R) Xeon(R) Gold 6248 CPU @ 2.50GHz and all files are stored on HDD disks.

| Probes | Index size | Indexing time | Indexing memory |
| --- | --- | --- | --- |
| 10,000 | 955 GB | 10 h 11 m | 69.77 GB |
| 20,000 | 972 GB | 12 h 11 m | 77.03 GB |
| 40,000 | 1,029 GB | 19 h 50 m | 86.59 GB |

**Table. S2.** Search performance on GTDB r214 complete dataset on three queries with different numbers of probes. We used 3 queries. Gene SecY (termed a rare gene), a 16s rRNA gene, and a plasmid (accessions are available in Methods). We report the number of genomes with alignments (genome his), and how many of these matched specific criteria. QcovHSP is the query coverage per High-scoring Segment Pair (HSP). Pident is the percentage identity. LexicMap `search` command used default parameters. The computation environment was the same as Table S1.

| Query (length) | Probes | Search time | Search memory | Genome hits | Genome hits (qcovHSP≥70) | Genome hits (qcovHSP≥70 and pident≥80) |
| --- | --- | --- | --- | --- | --- | --- |
| **A rare gene**<br>1,299 bp | 10,000 | 67 s | 1.77 GB | 6,100 | 6,068 | 2,355 |
|  | 20,000 | 70 s | 2.42 GB | 6,230 | 6,197 | 2,356 |
|  | 40,000 | 64 s | 3.96 GB | 5,882 | 5,848 | 2,356 |
| **A 16S rRNA gene**<br>1,542 bp | 10,000 | 749 s | 3.35 GB | 303,037 | 227,095 | 117,104 |
|  | 20,000 | 723 s | 4.31 GB | 303,664 | 227,275 | 117,105 |
|  | 40,000 | 714 s | 6.14 GB | 305,360 | 227,264 | 117,107 |
| **A plasmid**<br>52,830 bp | 10,000 | 483 s | 4.31 GB | 60,800 | 755 | 755 |
|  | 20,000 | 627 s | 5.86 GB | 63,517 | 1,191 | 1,191 |
|  | 40,000 | 732 s | 7.52 GB | 64,970 | 1,191 | 1,191 |

**Table. S3.**
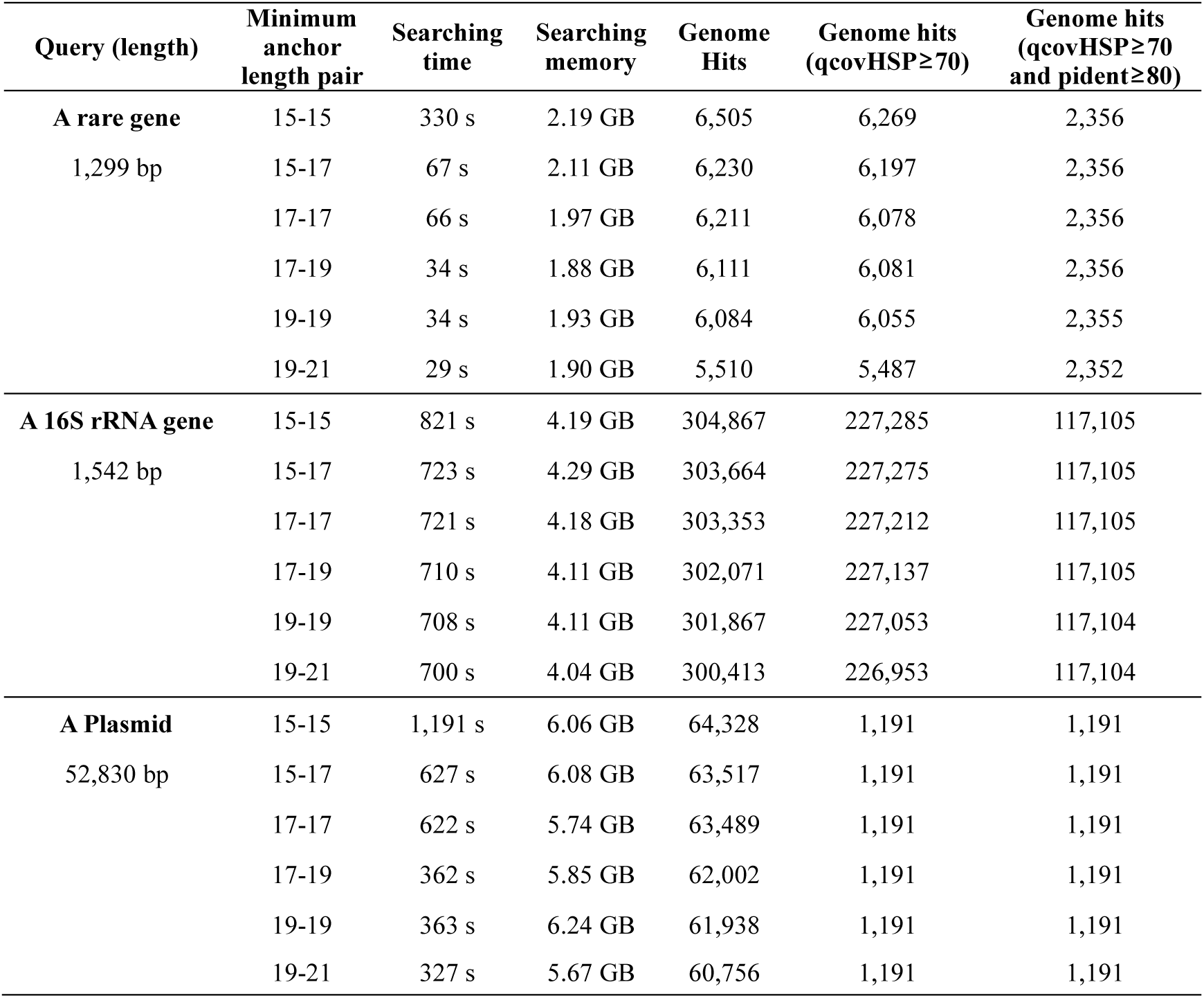
Searching performance on three queries with different paired minimum anchor length thresholds. The index was built with 20,000 probes from 402,538 prokaryotic assemblies from GTDB r214 complete dataset. LexicMap `search` command used default parameters except for the paired minimum anchor length thresholds (*P* and *P’*, see methods). The computation environment was the same as Table S1.

| Query (length) | Minimum anchor length pair | Searching time | Searching memory | Genome Hits | Genome hits (qcovHSP≥70) | Genome hits (qcovHSP≥70 and pident≥80) |
| --- | --- | --- | --- | --- | --- | --- |
| **A rare gene**<br>1,299 bp | 15-15 | 330 s | 2.19 GB | 6,505 | 6,269 | 2,356 |
|  | 15-17 | 67 s | 2.11 GB | 6,230 | 6,197 | 2,356 |
|  | 17-17 | 66 s | 1.97 GB | 6,211 | 6,078 | 2,356 |
|  | 17-19 | 34 s | 1.88 GB | 6,111 | 6,081 | 2,356 |
|  | 19-19 | 34 s | 1.93 GB | 6,084 | 6,055 | 2,355 |
|  | 19-21 | 29 s | 1.90 GB | 5,510 | 5,487 | 2,352 |
| **A 16S rRNA gene**<br>1,542 bp | 15-15 | 821 s | 4.19 GB | 304,867 | 227,285 | 117,105 |
|  | 15-17 | 723 s | 4.29 GB | 303,664 | 227,275 | 117,105 |
|  | 17-17 | 721 s | 4.18 GB | 303,353 | 227,212 | 117,105 |
|  | 17-19 | 710 s | 4.11 GB | 302,071 | 227,137 | 117,105 |
|  | 19-19 | 708 s | 4.11 GB | 301,867 | 227,053 | 117,104 |
|  | 19-21 | 700 s | 4.04 GB | 300,413 | 226,953 | 117,104 |
| **A Plasmid**<br>52,830 bp | 15-15 | 1,191 s | 6.06 GB | 64,328 | 1,191 | 1,191 |
|  | 15-17 | 627 s | 6.08 GB | 63,517 | 1,191 | 1,191 |
|  | 17-17 | 622 s | 5.74 GB | 63,489 | 1,191 | 1,191 |
|  | 17-19 | 362 s | 5.85 GB | 62,002 | 1,191 | 1,191 |
|  | 19-19 | 363 s | 6.24 GB | 61,938 | 1,191 | 1,191 |
|  | 19-21 | 327 s | 5.67 GB | 60,756 | 1,191 | 1,191 |

**Table. S4.**
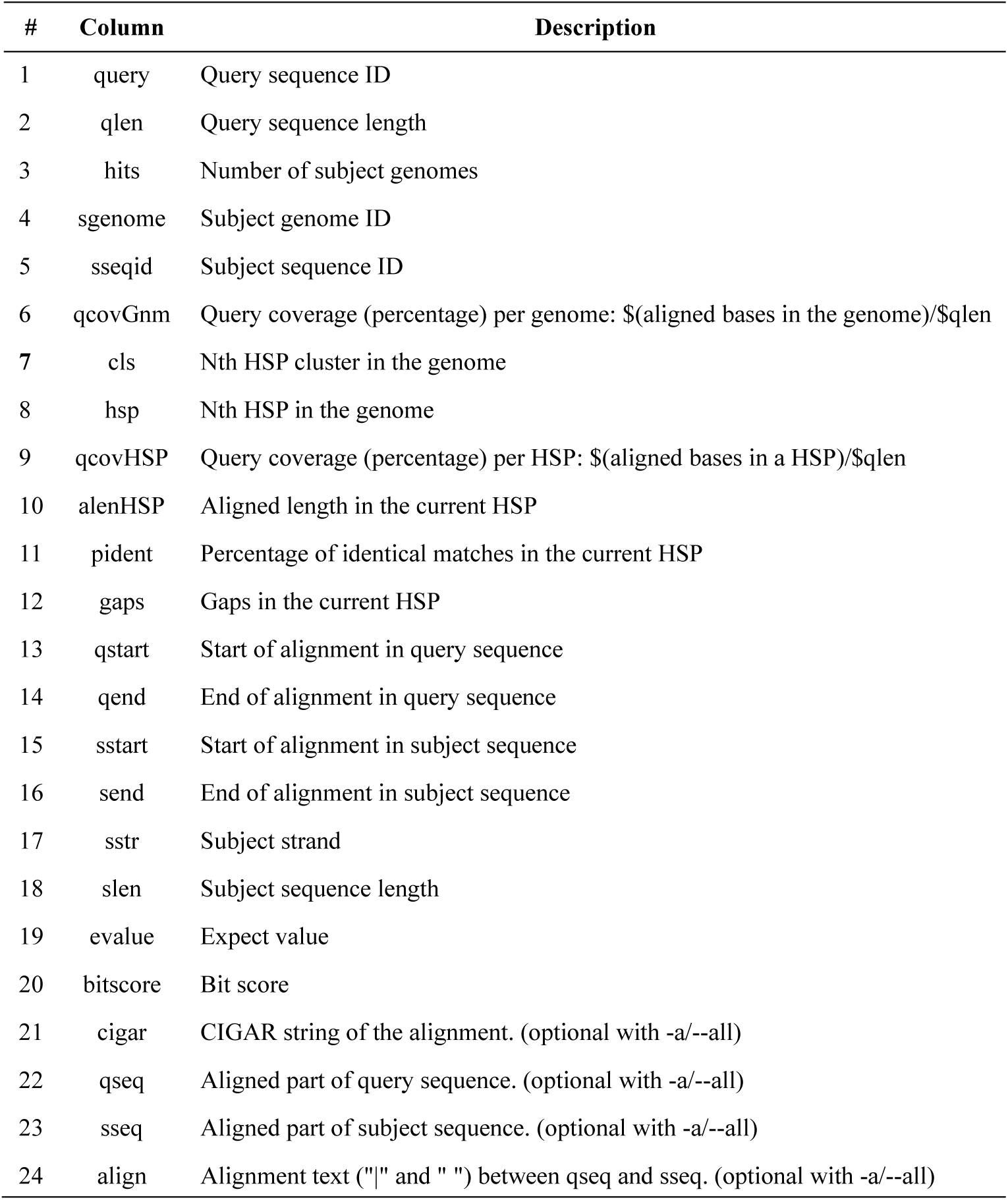
LexicMap alignment output format details.

**Table. S5.** Details of 10 bacterial genomes for the alignment rate comparison.

| Assembly | TaxID | Species | Contigs | Genome size |
| --- | --- | --- | --- | --- |
| GCF_000005845.2 | 562 | *Escherichia coli* | 1 | 4,641,652 |
| GCF_000240185.1 | 573 | *Klebsiella pneumoniae* | 7 | 5,682,322 |
| GCF_000013425.1 | 1280 | *Staphylococcus aureus* | 1 | 2,821,361 |
| GCF_000006945.2 | 28901 | *Salmonella enterica* | 2 | 4,951,383 |
| GCF_001457635.1 | 1313 | *Streptococcus pneumoniae* | 1 | 2,110,968 |
| GCF_000195955.2 | 1773 | *Mycobacterium tuberculosis* | 1 | 4,411,532 |
| GCF_000006765.1 | 287 | *Pseudomonas aeruginosa* | 1 | 6,264,404 |
| GCF_008632635.1 | 470 | *Acinetobacter baumannii* | 2 | 3,980,230 |
| GCF_018885085.1 | 1496 | *Clostridioides difficile* | 2 | 4,095,894 |
| GCF_022869705.1 | 1351 | *Enterococcus faecalis* | 1 | 2,866,948 |

**Table. S6.** Alignment performance of LexicMap on GenBank+RefSeq dataset. Sequence identifiers are available in Methods.

| Query | Query length | Hits (total) | Hits (high) | Hits (medi) | Hits (low) | Time | RAM |
| --- | --- | --- | --- | --- | --- | --- | --- |
| A rare gene | 1,299 bp | 41,718 | 11,746 | 115 | 29,857 | 3m:06s | 4.0 GB |
| A 16S rRNA gene | 1,542 bp | 1,955,167 | 245,884 | 501,691 | 1,207,592 | 32m:59s | 11.1 GB |
| A plasmid | 52,830 bp | 560,330 | 96 | 15,370 | 544,864 | 52m:22s | 14.5 GB |
| 1033 AMR genes | 1 kb (median) | 30,967,882 | 7,636,386 | 4,858,063 | 18,473,433 | 15h:52m:08s | 24.9 GB |

**Fig. S1.**
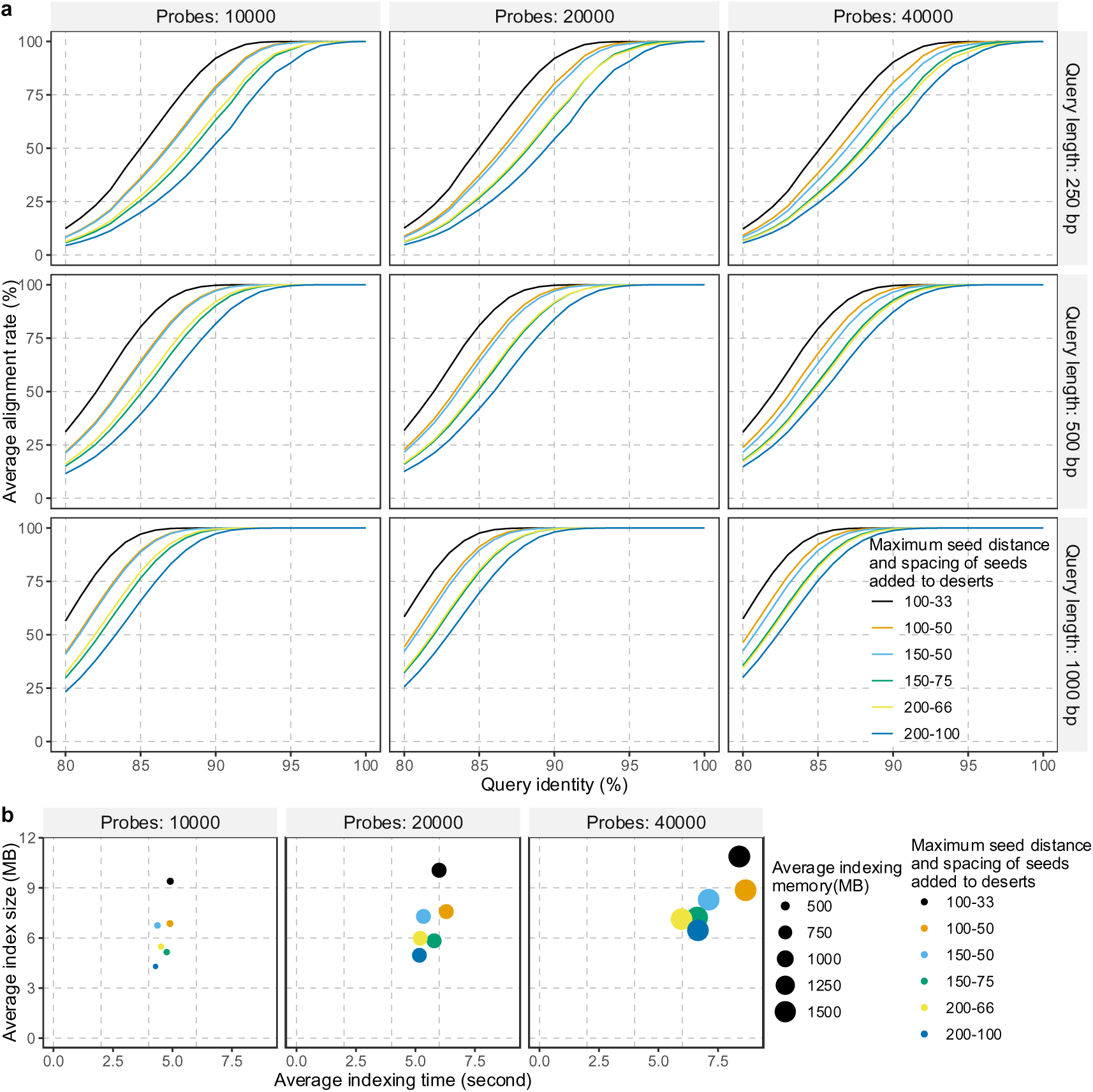
Performance of LexicMap with different numbers of probes and seed densities. **a)** Robustness to sequence divergence. **b)** Average index sizes of single genomes and indexing performances. The same 10 reference genomes and simulated queries are the same as those in Fig.3.

**Fig. S2.**
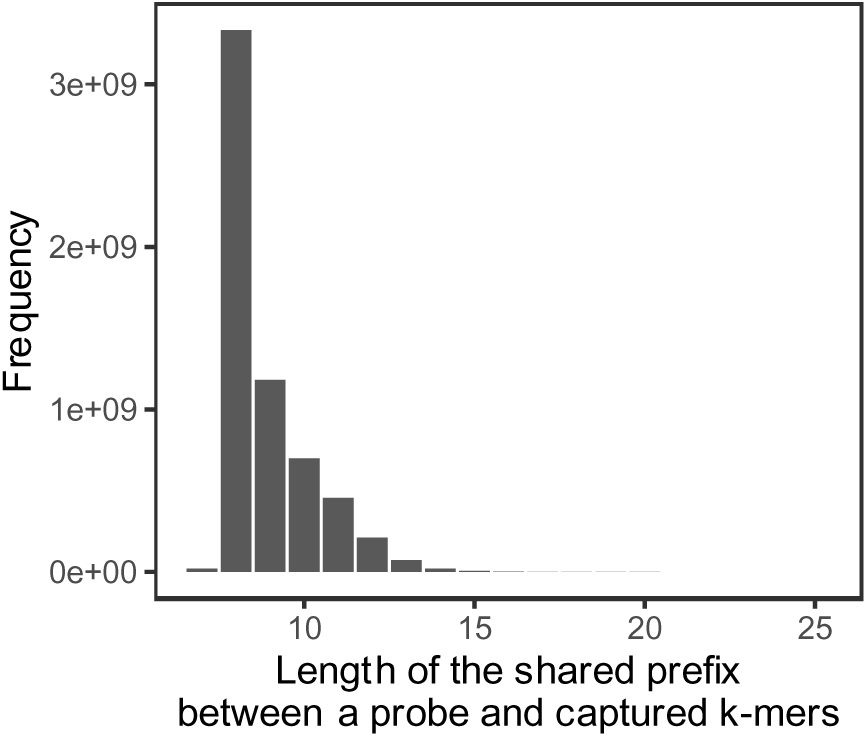
Histogram of shared prefix lengths between probes and the captured *k*-mers. Using a LexicMap index of GTDB r214 representative genomes (n=85,205), we show the lengths of shared prefixes between the 20,000 fixed probes and the captured *k*-mers..

**Fig. S3.**
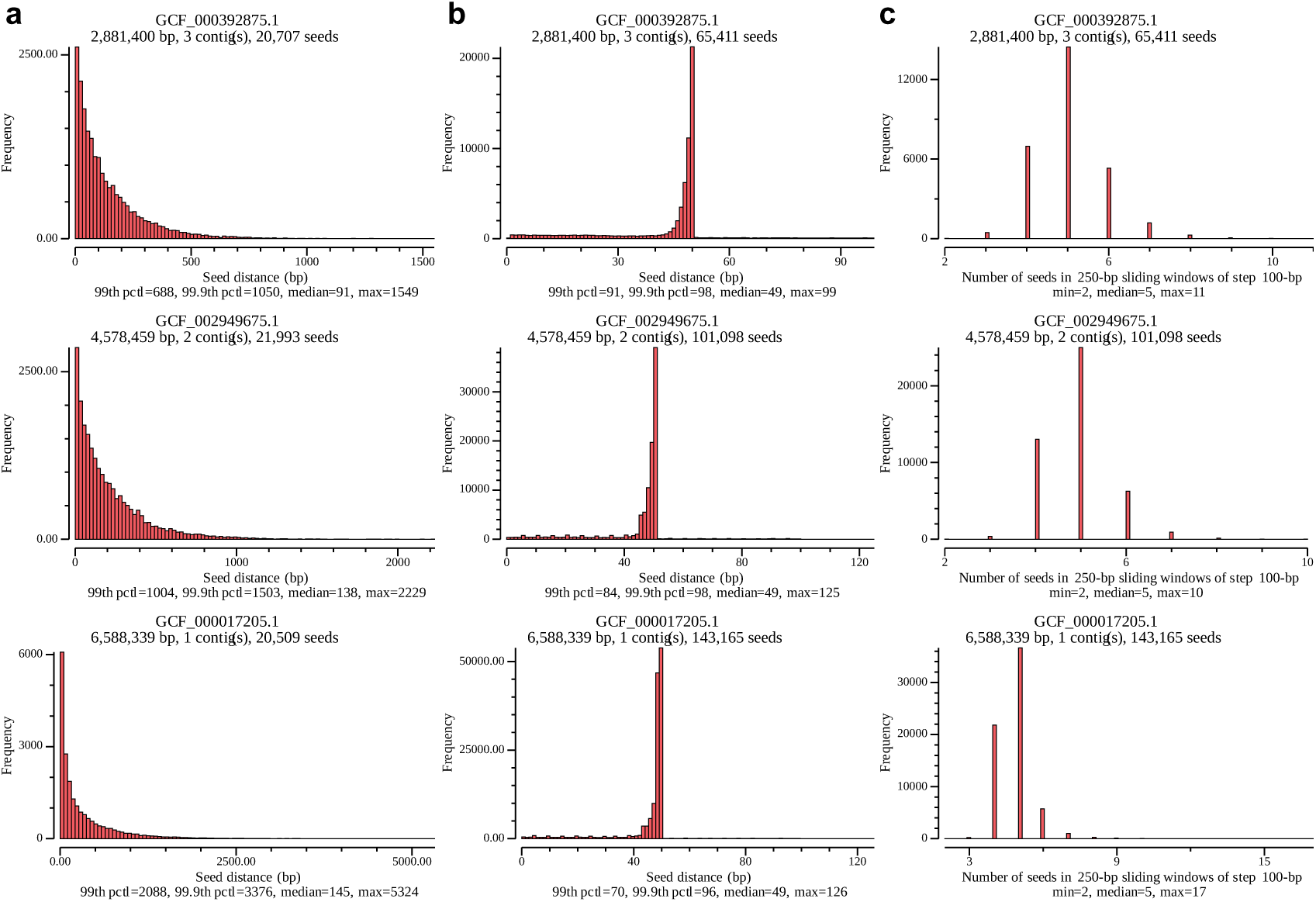
Histograms of seed distance and seed number in sliding windows. Three genomes of different genome sizes are used for illustration. a) Histograms of seed distance before filling seed deserts. b) Histograms of seed distance after filling deserts. Note that in assemblies GCF_002949675.1 and GCF_000017205.1, the two maximum seed distances larger than 100 bp resulted from low-complexity regions. c) Histograms of seed number in 250-bp sliding windows after filling deserts.

**Fig. S4.**
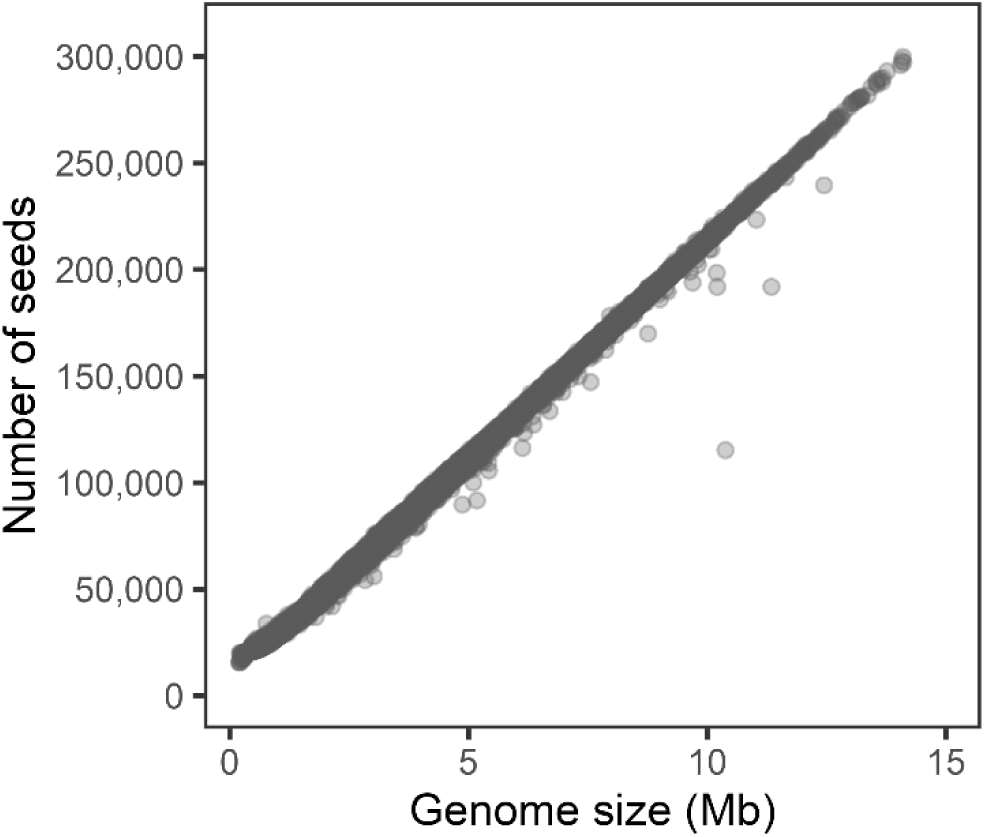
Number of seeds vs genome size. A LexicMap index was built from 85,205 GTDB r214 representative genomes with default parameters and “--save-seed-pos". The plot shows the number of seeds in each genome, versus the genome size (number of bases in Megabases).

**Fig. S5.**
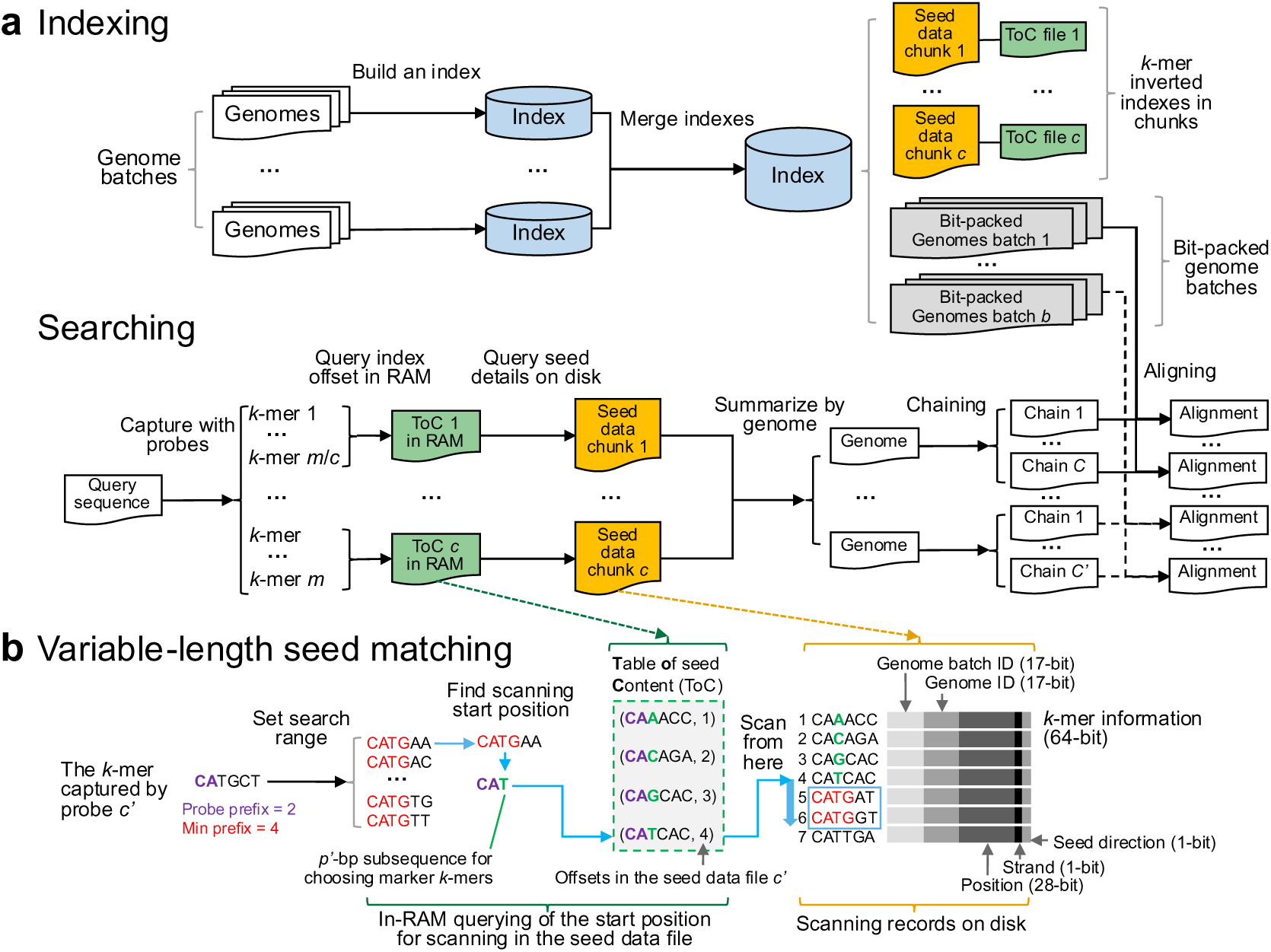
Architecture of LexicMap. a) Indexing and searching workflow. b) Variable-length seed matching.

**Fig. S6.**
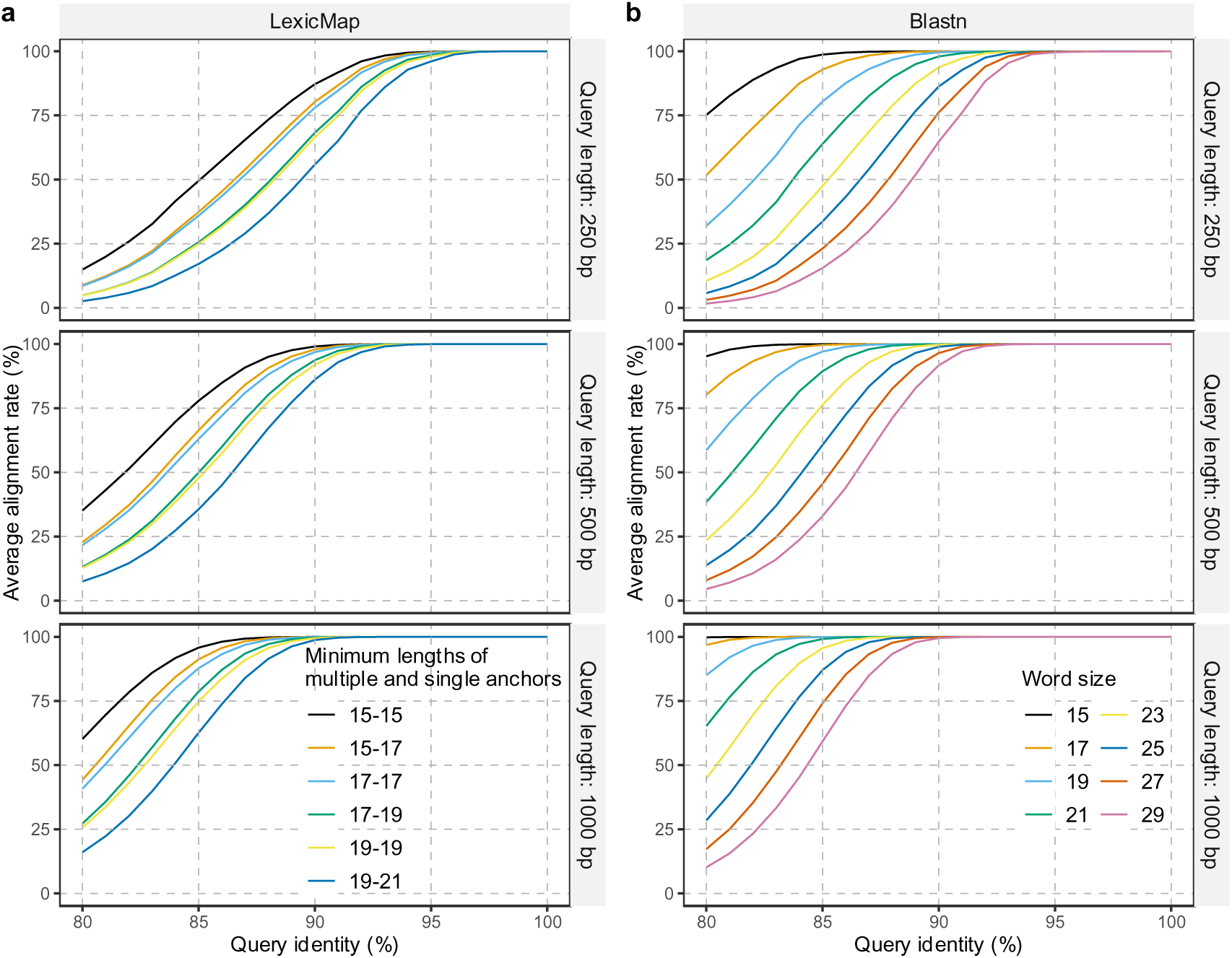
Robustness of LexicMap and Blastn to sequence divergence. LexicMap uses different paired minimum anchor length thresholds, and Blastn uses a range of word sizes. The 10 reference genomes and simulated queries are the same as those in Fig.3.

**Fig. S7.**
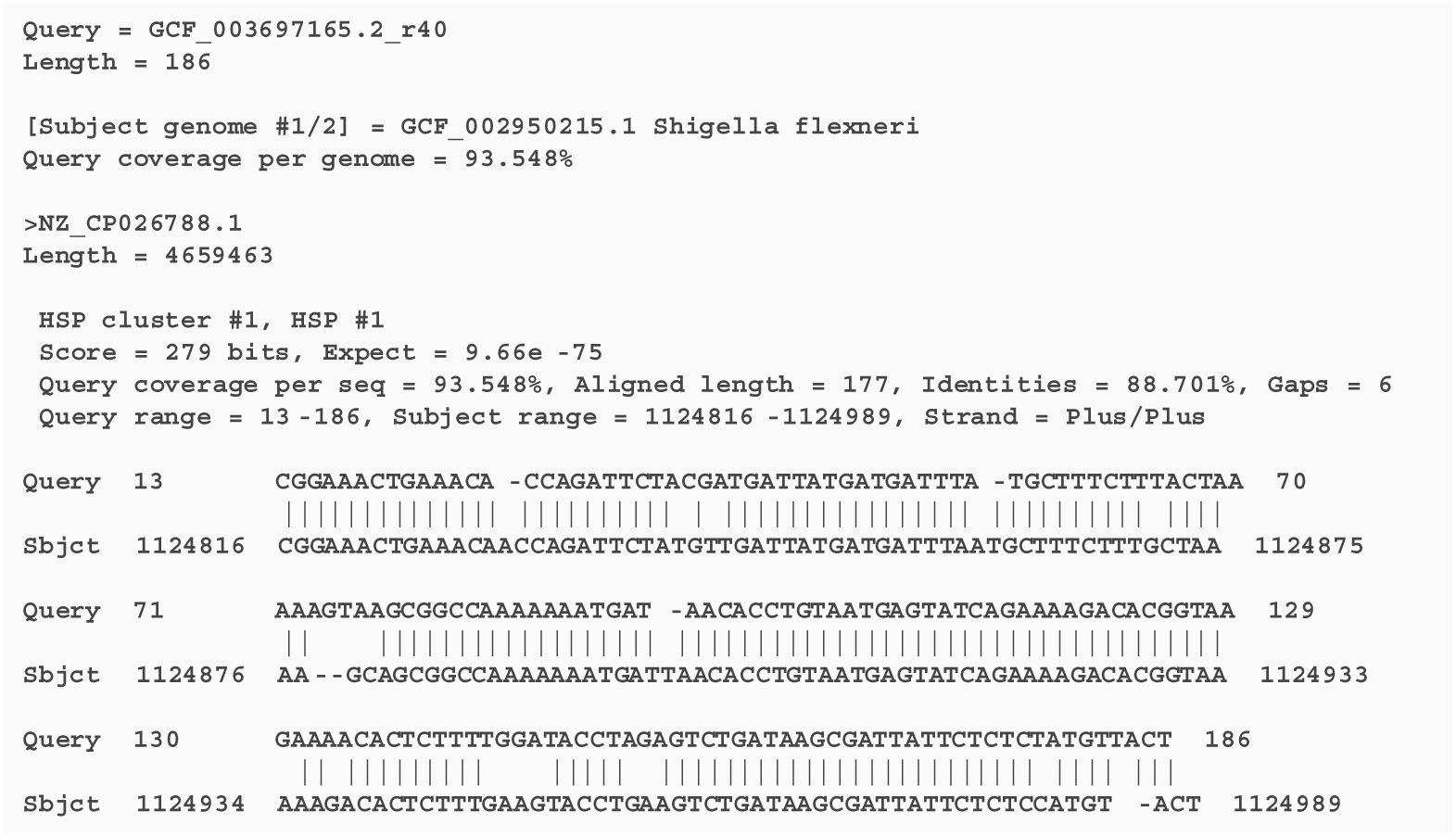
Blast-style pairwise output format in LexicMap. Note that, in addition to data that BLAST outputs, LexicMap also provides subject genome information (here *Shigella flexneri*) and query coverage per genome.

**Fig. S8.**
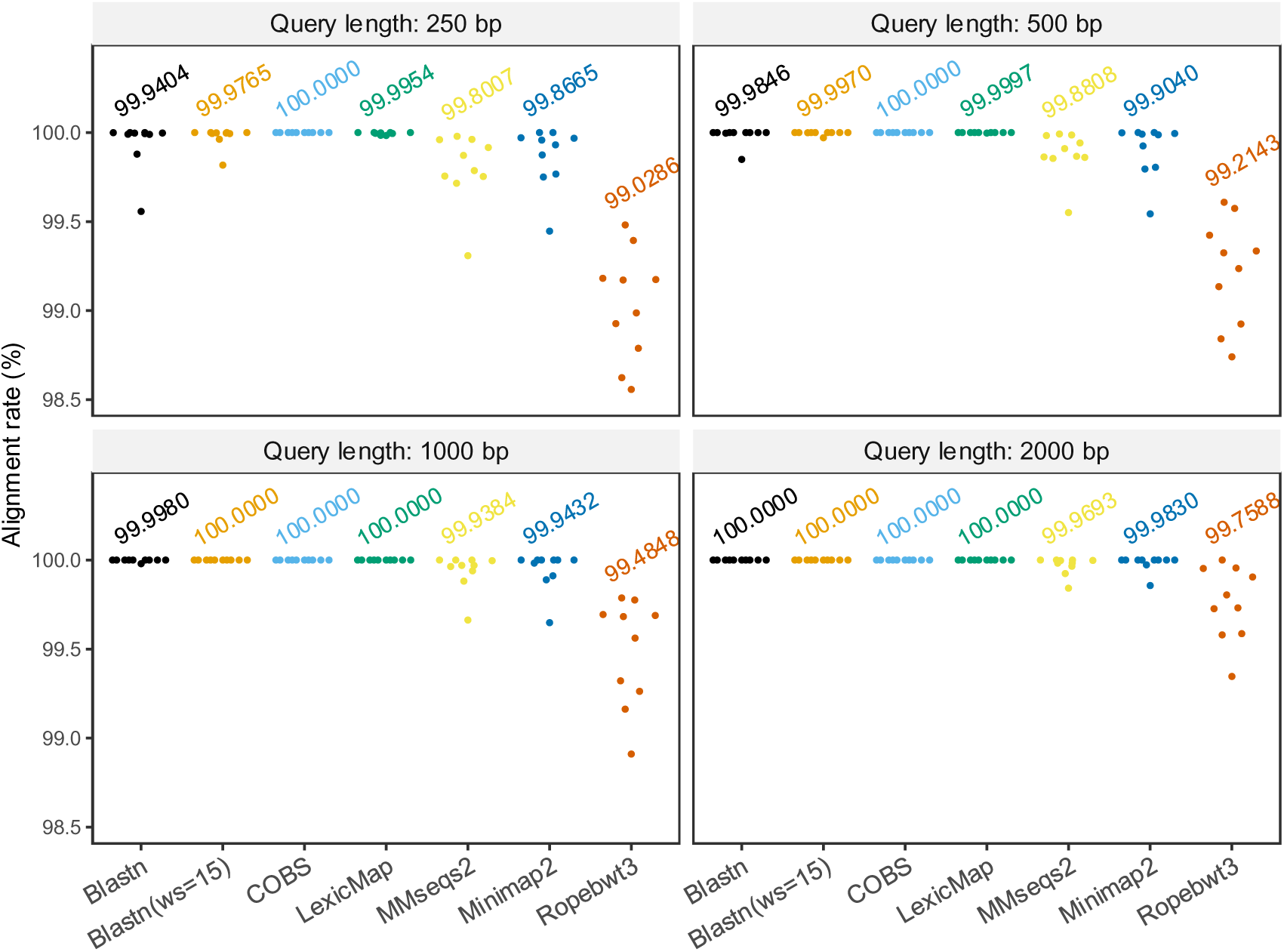
Alignment rates of simulated mutation-free queries. Queries with 100% identity (a subset of the experiment shown in in Fig.3) are aligned; the proportion of these which are aligned to the right place are shown here. Text labels are the mean alignment rates.

